# Kinetochore mutations and histone phosphorylation pattern changes preceded holo- and macro-monocentromere evolution

**DOI:** 10.1101/2025.05.06.652342

**Authors:** Yi-Tzu Kuo, Pavel Neumann, Jianyong Chen, Jörg Fuchs, Veit Schubert, Katrin Kumke, Mariela Analia Sader, Michael Melzer, Zihao Zhu, Axel Himmelbach, Heiko Hentrich, Jiří Macas, Andreas Houben

**Affiliations:** Leibniz Institute of Plant Genetics and Crop Plant Research (IPK) Gatersleben, Corrensstrasse 3, 06466 Seeland, Germany; Biology Centre, Czech Academy of Sciences, Institute of Plant Molecular Biology, Branišovská 31, České Budějovice, CZ-37005, Czech Republic; Multidisciplinary Institute of Plant Biology, National Council for Scientific and Technical Research (CONICET)-National University of Córdoba, Córdoba, Argentina; Molecular Cell Biology, Joseph Gottlieb Kölreuter Institute for Plant Sciences (JKIP), Karlsruhe Institute of Technology, Fritz-Haber-Weg 4, 76131 Karlsruhe, Germany

**Keywords:** divergent evolution, holocentromere, genome synteny, kinetochore composition, centromere plasticity

## Abstract

Centromeres are essential for kinetochore assembly and spindle attachment. While chromosomes of most species are monocentric with a single centromere, a minority exhibit holocentricity, with a centromere along the chromatid length. Sporadic emergence of holocentricity suggests multiple independent transitions. To explore this, we compared the centromere and (epi)genome organization of two sister genera with contrasting centromere types: *Chamaelirium luteum* with large “macro-monocentromeres” and *Chionographis japonica* with holocentromeres. Both exhibit chromosome-wide histone phosphorylation patterns distinct from typical monocentric species. Kinetochore analysis revealed similar chimeric *Borealin* in both species, with additional *KNL2* loss and *NSL1* chimerism in *Cha. luteum*. The broad-scale synteny between both genomes supports *de novo* holocentromere formation in *Chi. japonica*. Despite sharing features with both centromere types, macro-monocentromeres do not represent a direct link between mono- and holocentromeres. We propose a model for the divergent evolution involving kinetochore gene mutations, altered histone phosphorylation patterns, and centromeric satellite DNA amplification.

## Introduction

Centromeres – the constricted regions of chromosomes that connect sister chromatids – are essential for chromosome segregation in eukaryotes. Most organisms harbor monocentric chromosomes, which are characterized by primary constrictions ^1^, where the kinetochore assembles and spindle microtubules attach. A minority of species harbor an atypical centromeric organization known as holocentricity, in which centromeres are distributed over the entire chromatid length. Due to the telomere-to-telomere distribution of the holocentromere, sister chromatids cohere throughout their entire lengths and appear in mitotic chromosomes as two parallel structures, without a primary constriction ^2^. Each holocentromere comprises multiple ‘centromere units’, which, depending on the species investigated, possess either centromere-specific or non-specific DNA and on which a functional kinetochore protein complex is formed ^3^. In contrast, monocentromeres are composed of a single centromere unit per chromosome, although also monocentrics with a few neighboring centromere units per chromosome have been observed, e.g., in *Pisum* and *Lathyrus* ^4,5^ and the beetle *Tribolium castaneum* ^6^, forming so-called ‘meta-polycentric’ chromosomes ^5^.

The centromere-specific histone H3 (CENH3, also known as CENPA) specifies centromere identity in most species. It serves as a platform for kinetochore assembly, which includes the inner kinetochore constitutive centromere associated network (CCAN) and the outer kinetochore KMN complex (KNL1c, MIS12c, and NDC80c) for spindle microtubules ^7^. The precise balance between centromere-to-spindle-based tension and sister chromatid cohesion ensures accurate chromosome segregation ^8^, monitored by the spindle assembly checkpoint (SAC), which verifies proper chromosome-spindle attachments before anaphase. Cell cycle-dependent histone phosphorylation is crucial for recruiting SAC proteins in this process ^9^.

Histone H3 phosphorylation of the pericentromeric region is part of the histone modification network acting to control centromere function. These modifications can create a permissive environment for the correct assembly and function of centromeres ^9^. In plants, with the onset of mitosis, only pericentromeres, where sister chromatids cohere, undergo histone H3 phosphorylation at serine 10 and 28 ^10,11^. In holocentrics, due to their distinct centromere arrangement, these epigenetic marks are dispersed along the whole length of condensed chromosomes. In monocentric species, meanwhile, both epimarks are enriched adjacent to primary constrictions ^11^. Histone H3 threonine 3 phosphorylation originates at pericentromeres in prophase and evenly distributes along chromosome arms at prometaphase in monocentric chromosomes ^12,13^. In contrast, histone H2A threonine 120 (H2AT120ph; the threonine position refers to human H2A, and the corresponding positions differ between species) phosphorylation marks mirror the distribution of CENH3 in both centromere types ^14,15^.

Since holocentric species appear sporadically within phylogenetic lineages that have monocentric chromosomes, it is believed that holocentric chromosomes evolved from monocentric ones. Such a one-way transition happened multiple times in various green algae, protozoans, invertebrates, and different plant families ^16^. As a consequence of their independent evolution, holocentromeres are diverse in composition and organization (reviewed in ^3,17,18^). However, the mechanisms underlying this mono- to holocentromere transition are not yet understood. Analyzing closely related species with contrasting centromere types might illuminate this process. In the parasitic plant genus *Cuscuta*, the mono- to holocentromere transition is associated with extensive changes in genes responsible for the structure and regulation of the kinetochore ^19^. In insects such as Lepidoptera, the transition to holocentricity is related to the loss of CENH3 ^20^, leading to a permissive chromatin state-based centromere identity ^21^ and to CCAN-mediated kinetochore assembly in holocentric taxa ^22^.

An independent transition to holocentricity occurred in the plant tribe Chionographideae, constituting the holocentric genus *Chionographis* and the monocentric, monotypic sister genus *Chamaelirium*, in the monocot family Melanthiaceae ^23–28^ (Figures 1A, S1A, and S1B). The two genera diverged about 23.5 million years ago (mya) and currently exhibit a disjunct distribution, with *Chionographis* found in East Asia and *Chamaelirium* in North America ^29^. It has been suggested that the two genera be reclassified as parts of the merged genus *Chamaelirium* due to their otherwise considerable morphological similarities ^30^.

**Figure 1.**
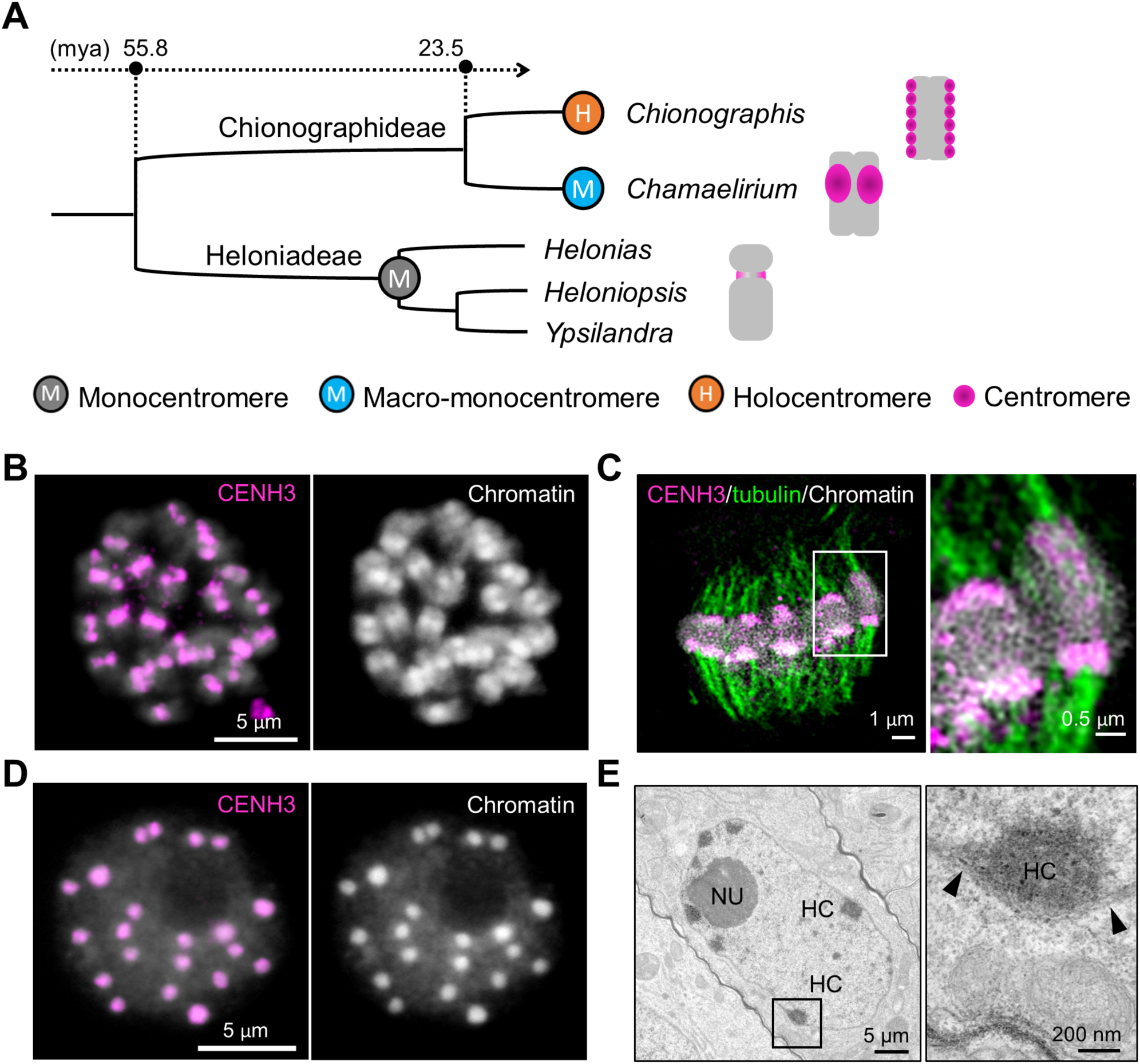
Primary constriction-free *Cha. luteum* chromosomes possess CENH3-positive macro-monocentromeres. (A) Phylogenetic tree of the tribes Chionographideae and Heloniadeae. Heloniadeae, including the genera *Helonias*, *Heloniopsis*, and *Ypsilandra*, is closely related to Chionographideae, with the two tribes diverging ∼ 55 mya ^29^. The schemata on the right show the overall chromosome and centromere (magenta) structure. (B) Primary constriction-free metaphase chromosomes show extensive CENH3 immunosignals (magenta). (C) Microtubules (green) attach to the poleward surface of macro-monocentromeres as observed by super-resolution microscopy (3D-SIM). The enlargement shows the colocalization between CENH3 (magenta) and microtubules (green). (D) CENH3 signals cluster in brightly DAPI-stained chromocenters of an interphase nucleus. (E) Transmission electron micrograph of a *Cha. luteum* interphase nucleus. Electron-dense heterochromatic chromocenters (HC) are often located in proximity to the double-layered nuclear membrane (further enlarged insert, arrows). NU, nucleolus. Chromatin was counterstained with DAPI.

Notably, the monocentric chromosome-typical primary constriction is absent in *Chamaelirium luteum* ^28^. Instead, the species features unusually large heterochromatic centromeres that protrude poleward at metaphase (Figure S1A). It was assumed that the ‘macro-monocentromeres’ of *Cha. luteum* might represent a precursor state from which holocentromeres in the sister genus *Chionographis* evolved ^28^. In the holocentric *Chi. japonica*, each holocentromere is composed of merely 7 to 11, ∼1.9 Mb-sized, minisatellite DNA-based, CENH3-positive centromere units that are arranged in a line along the poleward surface of each chromatid ^31^ (Figure S1B). The centromere units of *Chi. japonica* are up to 200 times larger than those described for the repeat-based holocentromeres in *Rhynchospora* species ^32^.

To gain insight into the evolution of atypical centromeres accompanying the mono- to holocentric transition, we resolved the genome and centromere organization of *Cha. luteum*. We compared the (epi)genome and kinetochore composition, as well as the genome synteny, taking advantage of the closely related species *Chi. japonica*, which possesses a contrasting centromere type and the same number of chromosomes (2*n*=24). Interestingly, the large macro-monocentromeres (up to 15 Mb) of *Cha. luteum* have characteristics in common with mono- and holocentromeres, but, in contrast to initial expectation, they do not constitute a direct link between mono- and holocentromeres. Instead, our comparative analysis of kinetochore composition revealed a loss of the *KNL2* gene and a chimeric origin of the *NSL1* (a MIS12c component) gene in *Cha. luteum* and of the *Borealin* gene in *Cha. luteum* and *Chi. japonica*. The observed conservation of genome synteny between the two species suggests that the holocentromere formation occurred *de novo* in *Chionographis*. Additionally, comparing both species to representatives of the closely related monocentric tribe Heloniadeae revealed that the cell cycle-dependent H3 phosphorylation along sister chromatid cohesion sites is required, but not sufficient for the formation of holocentromeres. Based on our findings, we discuss possible mechanisms driving the parallel transition from a typical monocentromere to either atypical macro-mono- or even holocentromeres.

## Results

### *Cha. luteum* chromosomes are monocentric despite the lack of a primary constriction

The primary constriction typical for monocentromere is absent in *Cha. luteum* chromosomes. Instead, large heterochromatic regions protrude on metaphase chromosomes (Figure S1B). To determine whether the so-called ‘macro-centromere’ ^28^ acts as an active centromere, a *Cha. luteum*-specific CENH3 antibody was generated as a marker. For this purpose, we analyzed the *Cha. luteum* transcriptome and identified a single *CENH3* gene. Its protein sequence phylogenetically grouped with that of the holocentric *Chi. japonica* (Figure S1D). As expected for CENH3 ^33^, amino acid sequences differed most at the N-terminal tail (Figure S1E). The first twenty N-terminal amino acids were used to generate the anti-CENH3 antibody. Immunostaining of somatic metaphase cells revealed CENH3 signals decorating the protruding ‘macro-monocentromeres’ (Figure 1B), where multiple spindle microtubules attach (Figure 1C and Video S1). During interphase, centromeres formed brightly DAPI-stained, almost equal-sized chromocenters (Figure 1D). The number of CENH3-positive chromocenters (17–24, on average 20, counted in 30 nuclei) is less than or equal to the number of chromosomes, as in the case of other chromocenter-forming monocentric species, such as *Arabidopsis thaliana* ^34^. A preferential distribution of chromocenters close to the nuclear double membrane was revealed (Figure 1E). A similar nuclear membrane preference of centromeric chromocenters was also found in the holocentric *Chi. japonica* as well as in *A. thaliana* ^31^. Our analysis of *Cha. luteum* demonstrates that the chromosome protrusion acts as a centromere and that a primary constriction is not a ubiquitous structural requirement for centromere function in a monocentric species.

### *Cha. luteum* exhibits exceptionally large satellite-based monocentromeres, ranking among the largest observed

The genome size of *Cha. luteum* was determined as 887.4 Mb/1C using flow cytometry. To resolve the sequence and size of the *Cha. luteum* ‘macro-monocentromeres’, we assembled its genome using a combination of HiFi reads with Hi-C scaffolding, resulting in a ∼745 Mb assembly representing ∼84% of the *Cha. luteum* genome. The refined assembly retained 92.7% complete BUSCOs across 649 contigs (N50=2.1 Mb, L50=87) (Table S1). Subsequent Hi-C scaffolding generated the top 22 scaffolds, each exceeding 10 Mb (10–68 Mb) used for downstream analysis (Table S2 and Figure S2A).

To determine whether the centromeres of *Cha. luteum* are composed of repetitive sequences, we first analyzed the repeat composition in its genome using RepeatExplorer and used the identified repeat clusters as references to assess their enrichment in CENH3-ChIP sequence data. Approximately 49% of the genome is composed of high-copy satellite repeats and transposable elements (Table S3). Among these, the 60-bp-monomer satellite cluster CL1, named ‘*Chama’*, is the most abundant repeat (9.3%) (Figure 2A). Its monomer sequence is rich in centromere-typical dyad symmetries, which may preferentially form a hairpin-loop structure (Figure 2A). FISH demonstrated centromeric positioning of the satellite cluster (Figure 2B), and its enrichment in the CENH3-precipitated fraction is further suggestive of a centromeric nature (Figure 2C). Additionally, the enrichment of CENH3-ChIP reads in the *Chama* satellite arrays of the assembly (Figure 2D), as well as the colocalization of CENH3-immuno and *Chama*-FISH signals in interphase chromocenters and heterochromatin regions of metaphase chromosomes (Figure 2E), confirmed its centromeric localization. In addition, the (AG/CT)_n_-containing CL51 and GC-rich CL176 repeat clusters, both with very low genome abundance (< 0.3%) and no sequence similarity to the *Chama* satellite, were also enriched in the CENH3-precipitated fraction (Figure 2C).

**Figure 2.**
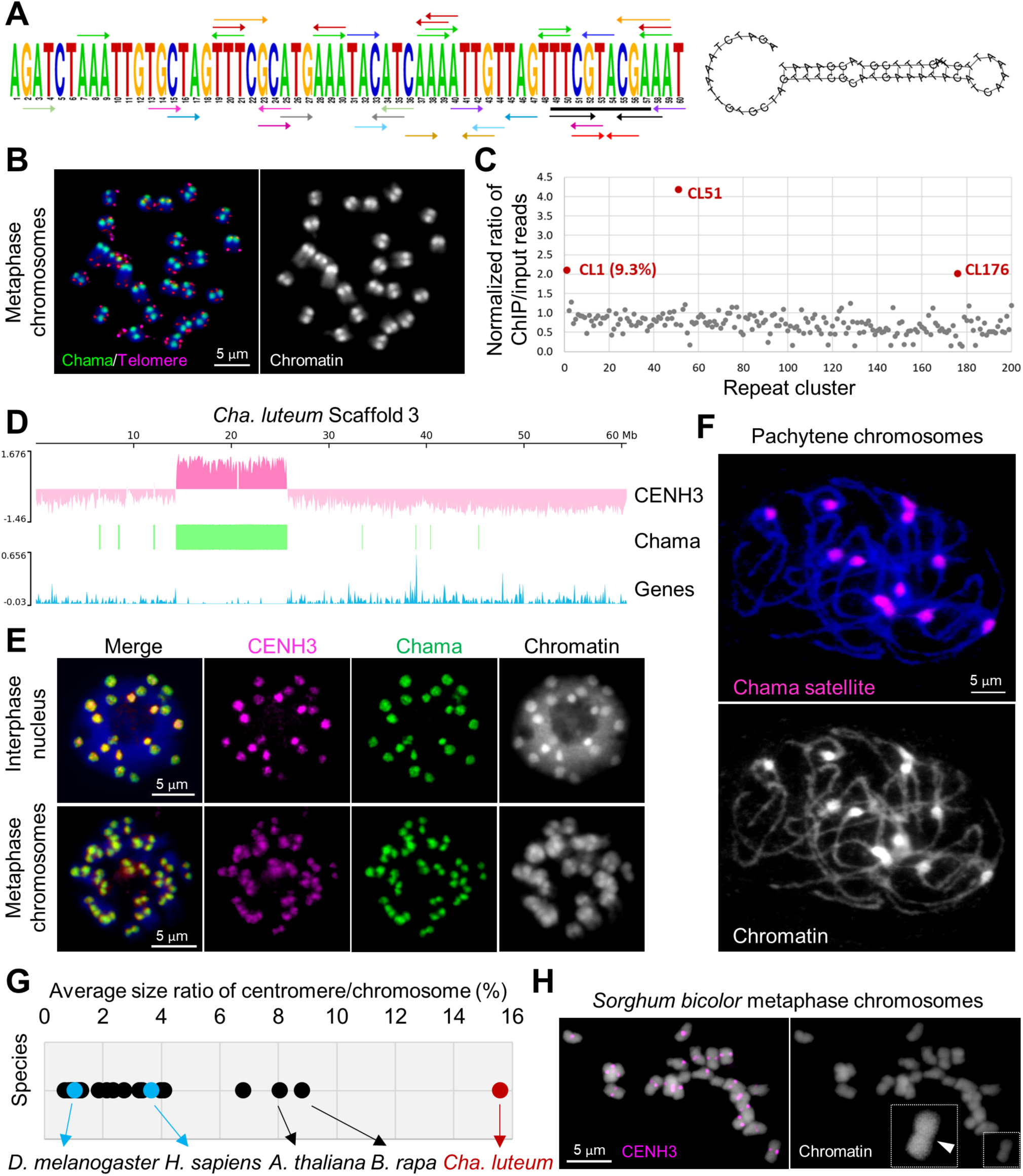
The macro-monocentromeres in *Cha. luteum* are satellite repeat-based. (A) The monomer sequence of the *Chama* satellite repeat. The colored arrows indicate dyad symmetries. The 9-bp TTCGTACGA (underlined in black) is shared between the 60-bp *Chama* monomer and the 23-bp *Chio* monomer sequences ^31^. Predicted hairpin loop structure formed by a *Chama* monomer. (B) Mitotic metaphase of *Cha. luteum* showing *Chama* (green) and telomere-specific (magenta) signals. (C) The genome proportion and normalized enrichment in CENH3-ChIPseq of the RepleatExplorer clusters. (D) Mapping of the CENH3-ChIPseq reads (pink), distribution of Chama satellite repeats (green) and of genic sequences (blue) to the 60 Mb-large scaffold 3. Note the strict enrichment of CENH3 on the Chama satellite array. (E) *Cha. luteum* interphase and metaphase chromosomes show colocalization of anti-CENH3 (magenta) and Chama repeat-specific (green) imunoFISH signals. (F) Chama satellite repeats locate in the knob-like structures of pachytene chromosomes. (G) The average size ratio between centromere and chromosome in monocentric species, including *Cha. luteum* (15.6%, red dot). The indicated species are *Arabidopsis thaliana* (8.1%), *Brassica rapa* (8.8%), *Homo sapiens* (3.7%, blue dot), and *Drosophila melanogaster* (1.1%, blue dot). The data of this plot are in Table S4. (H) The immunolabelling of CENH3 (magenta) in monocentric *Sorghum bicolor*. The chromosomal constriction in the enlargement is indicated by an arrowhead. Chromosomes were counterstained with DAPI.

Among the top 22 scaffolds of *Cha. luteum*, the eight CENH3-interacting *Chama* arrays range in size from 9.58 to 15.30 Mb, with an average size of 11.54 Mb (Table S2). The sequence arrays are highly homogeneous and consistent in their sequence orientation (Figure S2B), unlike the frequently alternating orientation of the centromeric *Chio* satellite arrays in *Chi. japonica* ^31^ (Figure S2C). Our FISH analysis of naturally extended pachytene chromosomes, which feature knob-like centromeres, in line with our sequencing results, ruled out the possibility that the macro-monocentromeres of *Cha. luteum* are formed of multiple adjacent centromere units typical for ‘meta-polycentric’ chromosomes, as the number of *Chama* signals equaled the number of chromosome bivalents (Figure 2F).

A comparison of CENH3-ChIPseq-defined centromere sizes, relative to chromosome sizes, across monocentric species showed that *Cha. luteum* has the proportionally largest centromeres reported to date (∼15.6% of an average chromosome) (Figure 2G and Table S4). For a direct comparison of the centromere size with a species possessing similar chromosome dimensions, we performed immunolabeling of *Sorghum bicolor* centromeres with a *S. bicolor*-specific CENH3 antibody, resulting in CENH3 signals at primary constrictions (Figure 2H) ^35^. Despite comparable chromosome sizes (*Cha. luteum* (887 Mb/1C, *n*=12) ≈ 73.9 Mb/chromosome, *S. bicolor* (789 Mb/1C, *n*=10) ≈ 78.9 Mb/chromosome), *Cha. luteum* exhibits substantially larger centromeres.

### The genomes of holocentric *Chi. japonica* and monocentric *Cha. luteum* share broad-scale synteny, with the exception of their centromeres

Centromeres play a crucial role in shaping genome architecture ^36^. To investigate whether the evolution of the two centromere types from the sister genera was accompanied by genome reshuffling, we analyzed syntenic orthologs of single-copy coding genes in the assembled scaffolds of *Cha. luteum* and the chromosome-level genome assembly of *Chi. japonica.* We identified 9,960 pairs of collinear genes, revealing large blocks and a high degree of genome conservation (Figures 3A and S3), althought their genomes diverged 23.5 million years ago ^29^. In particular, the chromosome-level arrangement of orthologs between chromosome 8 of *Chi. japonica* and scaffold 3 of *Cha. luteum* is identical (Figure 3B). The order and orientation of all six non-centromeric intervals of *Chi. japonica* chromosome 8 were conserved in the corresponding chromosome-sized scaffold 3 of *Cha. luteum.* Besides chromosome-sized syntenic regions, 11 large-scale inversions and four inter-chromosomal translocations (scaffolds 2, 6, 7, and 13 of *Cha. luteum*) were found (Figures 3A and S3). Nevertheless, local gene synteny was mainly conserved. Notably, syntenic blocks were split by centromere units in *Chi. japonica* (Figures 3B and S3), indicating *de novo* origin of centromere units in this holocentric species. In addition, ATAC-seq revealed that the entire *Cha. luteum* chromosome, except for the centromere, exists in an open chromatin state with an almost uniform distribution of genes (Figure 3B). This suggests that the *de novo* holocentromere formation sites *in Chi. japonica* correspond to gene-rich, open chromatin regions in the syntenic *Cha. luteum* genome. According to the synteny of centromere flanking regions between the two genomes, the positions of the monocentromeres in *Cha. luteum* (e.g., scaffold 3) and centromere units in *Chi. japonica* do not always correspond (Figures 3B and S3). Assuming that the centromere positions of *Cha. luteum* and of the ancestor of both species are conserved, the loss of monocentromeres was thus likely accompanied by *de novo* holocentromere formation in *Chionographis*.

**Figure 3.**
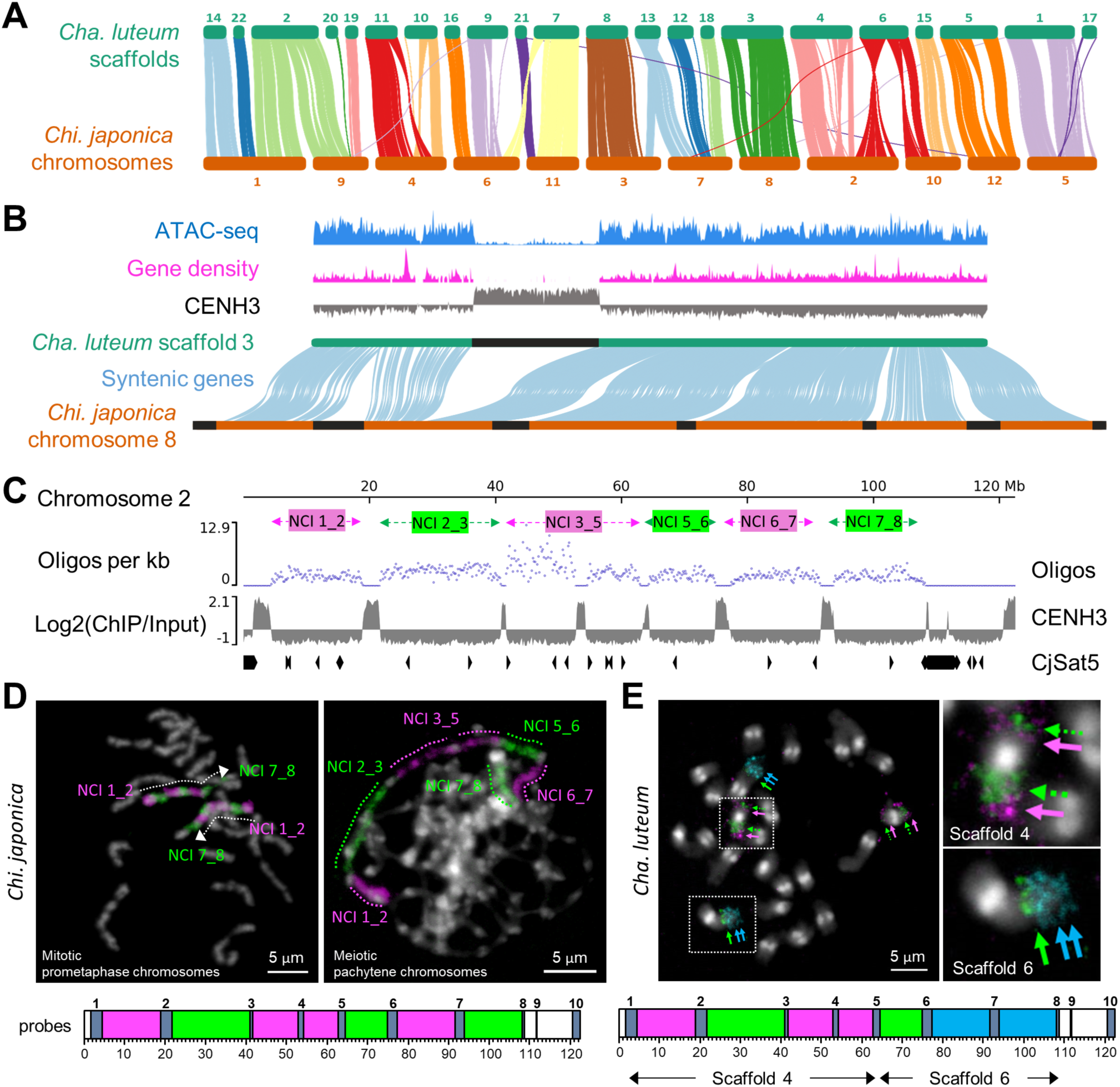
Holocentric *Chi. japonica* and monocentric *Cha. luteum* share broad-scale genome synteny, except for their centromeres. (A) Vizualization of syntenic orthologs of coding genes in the assembled scaffolds of *Cha. luteum* and the chromosome-level genome assembly of *Chi. japonica.* (B) Chromosome-level arrangement of orthologs between scaffold 3 of *Cha. luteum* and chromosome 8 of *Chi. japonica* is identical, with the order and orientation of all six non-centromeric intervals of *Chi. japonica* chromosome 8 conserved in scaffold 3 of *Cha. luteum.* The position of the monocentromere in *Cha. luteum* doesn’t correspond to a centromere unit in *Chi. japonica* according to the syntenic genes. The gene distribution (magenta) and ATAC-seq data (blue) of *Cha. luteum* show that the *de novo* centromere formation sites *in Chi. japonica* correspond to a high gene density and open chromatin state of the corresponding regions in the *Cha. luteum* genome. (C) Design of the oligo-FISH painting probes specific for six non-centromeric intervals (NCIs) between the first and the eighth centromere unit of *Chi. japonica* chromosome 2. (D) FISH mapping on mitotic prometaphase and meiotic pachytene chromosomes of *Chi. japonica* confirmed the accuracy of the sequence assembly and the specificity of these oligo-FISH probes. (E) In *Cha. luteum*, the *Chi. japonica* chromosome 2 based oligo-FISH signals were located on three arms of two chromosome pairs, corresponding to scaffolds 4 and 6, as predicted by sequence analysis (Figure S4). Chromosomes were counterstained with DAPI.

To confirm the sequence-deduced conservation of syntenic chromosomal blocks and *in silico*-identified rearrangements between the two species, we designed oligo-FISH painting probes specific for the six non-centromeric intervals (NCIs) between the first and the eighth centromere unit of *Chi. japonica* chromosome 2 (Figure 3C). FISH mapping on mitotic prometaphase and meiotic pachytene chromosomes of *Chi. japonica* confirmed the correctness of the sequence assembly and specificity of oligo-FISH probes (Figures 3D and S4A–C). In *Cha. luteum*, the signals of the six probes were located on three arms of two chromosome pairs (Figures 3E and S4A–C), corresponding to scaffolds 4 and 6, as predicted by our sequence analysis (Figures 3A, 3E, and S4D). Additionally, we mapped the sequences of all 12 *Chi. japonica*-based, chromosome-wide oligo pools to the top 22 scaffolds of *Cha. luteum*. This alternative *in silico* strategy verified the high chromosomal collinearity between the two genomes with contrasting centromere types as well as the absence of chromosome duplications (Figure S5A).

To determine whether this large-scale genome synteny is conserved beyond the tribe Chionographideae, we checked for a potential cross *in situ* hybridization in the closely related species of tribe Heloniadeae, which includes the genera *Helonias*, *Heloniopsis*, and *Ypsilandra*, and which diverged ∼55 mya from tribe Chionographideae (Figure 1A) ^29^. However, the same oligo-FISH painting probes revealed no detectable signals on the chromosomes of *Heloniopsis umbellata*. Moreover, comparative repeat analysis revealed no shared high-copy repeats between *H. umbellata* and *Cha. luteum* or *Chi. japonica* (Figure S5B). Thus, despite their contrasting centromere types, the genomes of *Cha. luteum* and *Chi. japonica* are highly syntenic, yet while being divergent from those of Heloniadeae species.

### Loss and alteration of kinetochore genes are associated with the transition to unconventional centromeres in Chionographideae

The protein composition of the kinetochore complex varies considerably across species with different centromere types ^18,19,37,38^. To determine whether the evolution of holo- and macro-monocentromeres in different members of the Chionographideae was driven by similar molecular mechanisms, we investigated the kinetochore composition of *Chi. japonica, Cha. luteum*, the related monocentric species *Heloniopsis orientalis* from tribe Heloniadeae (Figures 1A and S1C), and of *Phoenix dactylifera*, a phylogenetically distant monocot species for comparison. In total, we analyzed 29 structural and regulatory kinetochore proteins (Figure 4 and Table S5).

**Figure 4.**
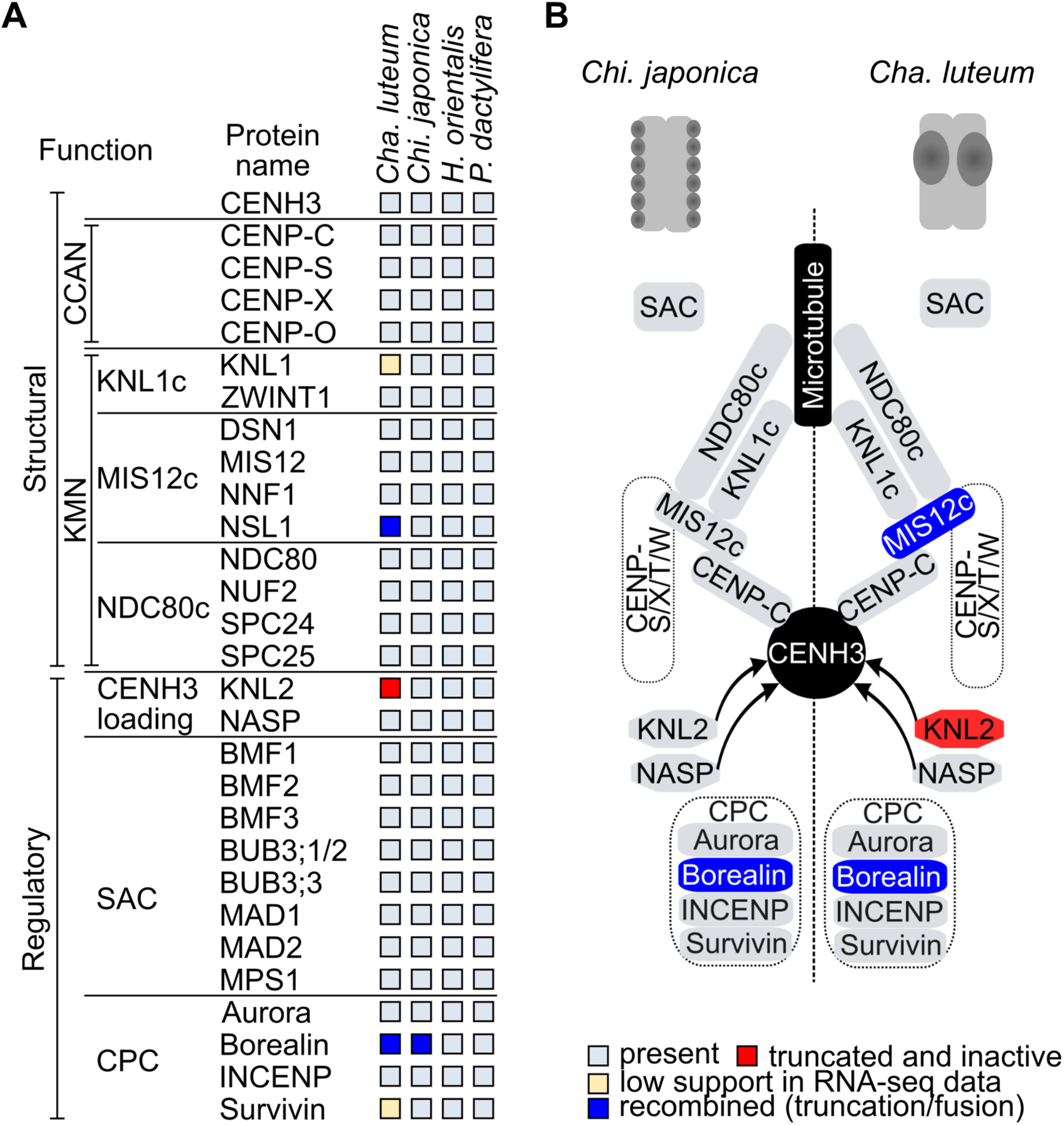
Changes in structural and regulatory kinetochore proteins in Chionographideae. (A) Comparative analysis of the kinetochore protein sequences in *Cha. luteum*, *Chi. japonica*, *Heloniopsis orientalis* and *Phoenix dactylifera.* The monocentric species *H. orientalis*, a species from the closely related tribe Heloniadeae, and *P. dactylifera*, a phylogenetically distant monocotyledon, were included for comparison. (B) Simplified schematic illustration of kinetochore structure and centromere organization in the chromosomes of *Chi. japonica* (left) and *Cha. luteum* (right). Proteins or complexes containing proteins that have changed or were lost in *Cha. luteum* or *Chi. japonica* are highlighted in blue or red. Sequence information of the analyzed kinetochore proteins are in Table S5. The figure was adapted from Neumann *et. al* (2023).

In this *in silico* analysis, we identified homologs for all tested kinetochore proteins in the transcriptomes of *Chi. japonica*, *H. orientalis*, and *P. dactylifera*. In *Cha. luteum*, we found no sequence similar to the CENH3-loading protein KNL2 (Figures 4A and 4B). A subsequent BLASTn search for the *KNL2* gene in the genome assembly of this species using the complete gene sequence of *Chi. japonica* ^31^, revealed only a short fragment at the 3’ end of the *KNL2* gene, spanning only two of the seven exons; notably, the exons encoding the conserved SANTA domain and CENPC-k motif of KNL2 were lost (Figure S6A). This non-functional *KNL2* fragment in *Cha. luteum* was found in the locus orthologous to the genomic region containing the functional *KNL2* gene in *Chi. japonica* (Figure S6B), indicating that it is a remnant of a formerly functional *KNL2* gene.

Comparative analysis of the identified kinetochore protein sequences revealed a remarkable N-terminal divergence in the MIS12c component NSL1 and the chromosomal passenger complex (CPC) module Borealin. Specifically, the N-terminus of NSL1 in *Cha. luteum* showed unique divergence (Figures 5A, S7A, and S7B). In contrast, Borealin sequences were similar between *Cha. luteum* and *Chi. japonica*, but the two were distinct from the highly conserved sequences in *H. orientalis* and *P. dactylifera* (Figures 5B and S7C). Domain analysis revealed that the divergent N-terminus of NSL1 in *Cha. luteum* shares similarity with the BLOC-1 Related Complex Subunit 6 (BORCS6) protein (Figure S7B), while the N-terminus of Borealin in *Cha. luteum* and *Chi. japonica* showed similarity to ribosomal protein S17 (RPS17) (Figure S7C). This suggests a chimeric origin of the *NSL1* gene in *Cha. luteum* and of the *Borealin* genes in both *Cha. luteum* and *Chi. japonica*, with the original N-terminus-coding regions replaced by sequences derived from *BORCS6* and *RPS17* genes, respectively (Figure 5). By identifying the donors of the N-terminal sequences, we determined the acquired N-terminal lengths to be 104 aa for NSL1 (59% of the protein) (Figure S7B) and 50 aa for Borealin (20% of the protein) (Figure S7C).

**Figure 5.**
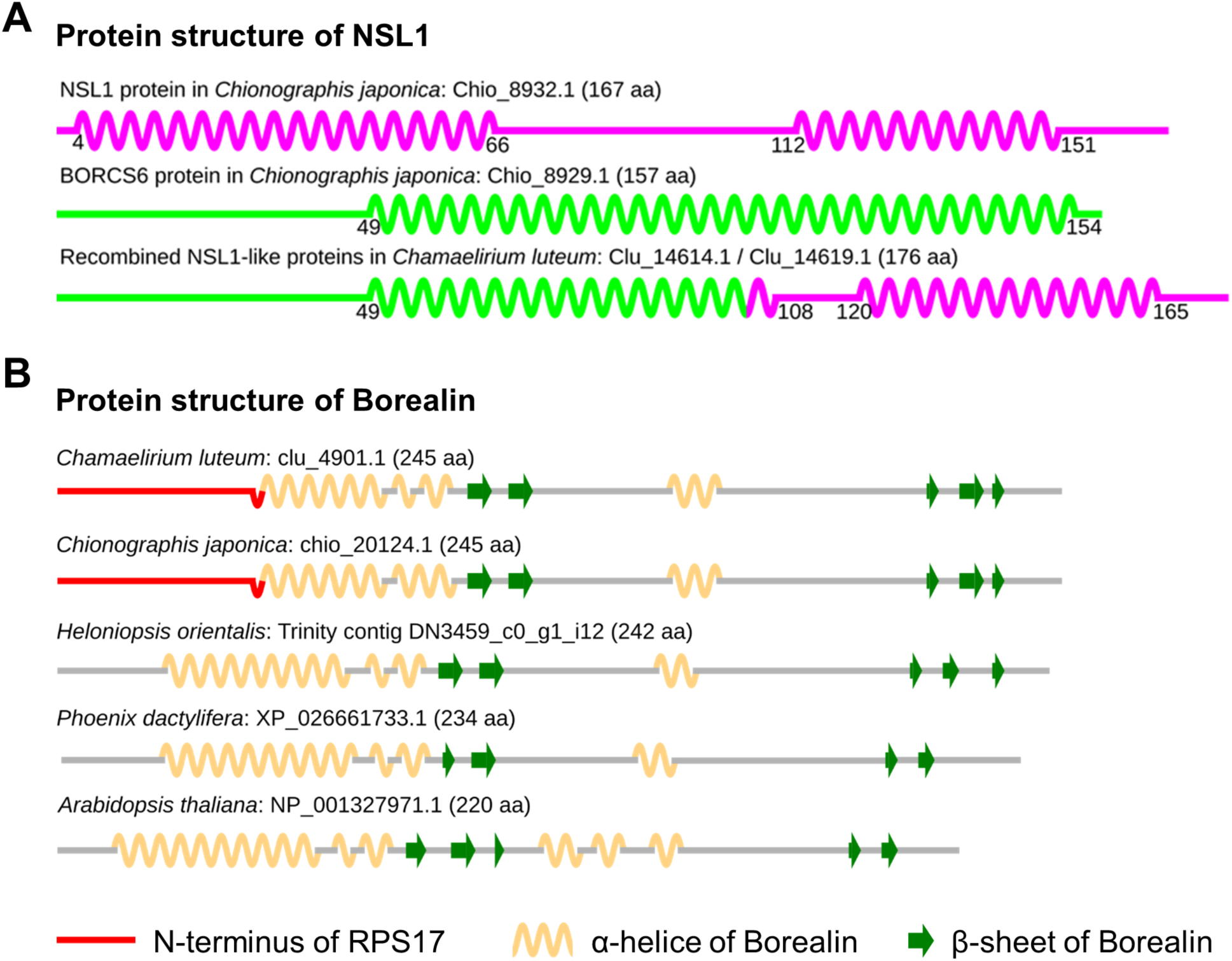
Chimeric origin of the *NSL1* and *Borealin* genes in *Cha. luteum*. (A) Comparison of the secondary protein structure of the intact and chimeric NSL1 proteins. NSL1 and BORCS6 proteins are shown in different colors to indicate the origin of the two parts in the chimeric NSL1. (B) Comparison of the secondary structure of Borealin proteins in three Melanthiaceae species, *Phoenix dactylifera*, and *Arabidopsis thaliana.* Although the structure is well conserved in all the species, the first α-helix is significantly shorter in *Cha. luteum* and *Chi. japonica*, which is due to the N-terminus of Borealin being replaced by the N-terminal fragment of ribosomal protein S17 (red). α-helices and β-sheets are shown as beige wavy lines and green arrows, respectively.

NSL1 is a structural component of the MIS12 complex, which consists of NSL1, MIS12, DSN1, and NNF1, and interacts with the other two outer kinetochore complexes (NDC80 and KNL1) via NSL1 and DSN1 as well as with the inner kinetochore proteins via DSN1 ^39,40^. Therefore, we wondered whether the observed change in NSL1 had an impact on kinetochore assembly in *Cha. luteum.* We took advantage of antibodies developed against MIS12 and NDC80 proteins of *Chi. japonica* as well as a highly versatile antibody against KNL1 in plants ^31,41^. All three antibodies specifically labeled holocentromeres in *C. japonica* and monocentromeres in Heloniadeae species ^31,41^. It was anticipated that the antibodies would mark the centromeres of *Cha. luteum* if the kinetochore composition remained unchanged because the sequences of the KNL1, MIS12, and NDC80 antibody target domains are identical between *Chi. japonica* and *Cha. luteum.* We failed to detect signals for KNL1 and NDC80 in *Cha. luteum*, even though we were able to successfully detect MIS12 using well-established immunodetection procedures (Figure S7D). Therefore, we hypothesize that the alteration of NSL1 either prevents KMN complex formation or that these two proteins are present at concentrations below the sensitivity of our immunostaining assay. The presence of the intact MIS12c component DSN1 likely explains why the centromeric recruitment of MIS12 appears to be unaltered.

### Monocentric *Cha. luteum* shows holocentromere-typical cell cycle-dependent histone phosphorylation patterns

Phosphorylation of histone H3 serves to prime chromatin for faithful chromosome segregation ^9^. Thus, we also analyzed the distribution of cell cycle-dependent, spindle assembly checkpoint-associated histone phosphorylation marks. Despite its monocentric chromosomes, anti-H3S10/S28/T3 phosphorylation signals were distributed throughout the mitotic metaphase chromosomes of *Cha. luteum* (Figure 6A), as in holocentric *Chi. japonica* ^31^ and other holocentric plants ^42^. Surprisingly, H2AT120 phosphorylation, an otherwise conserved cell cycle-dependent centromeric mark ^15,43^, was undetectable in both monocentric *Cha. luteum* and holocentric *Chi. japonica* ^31^.

**Figure 6.**
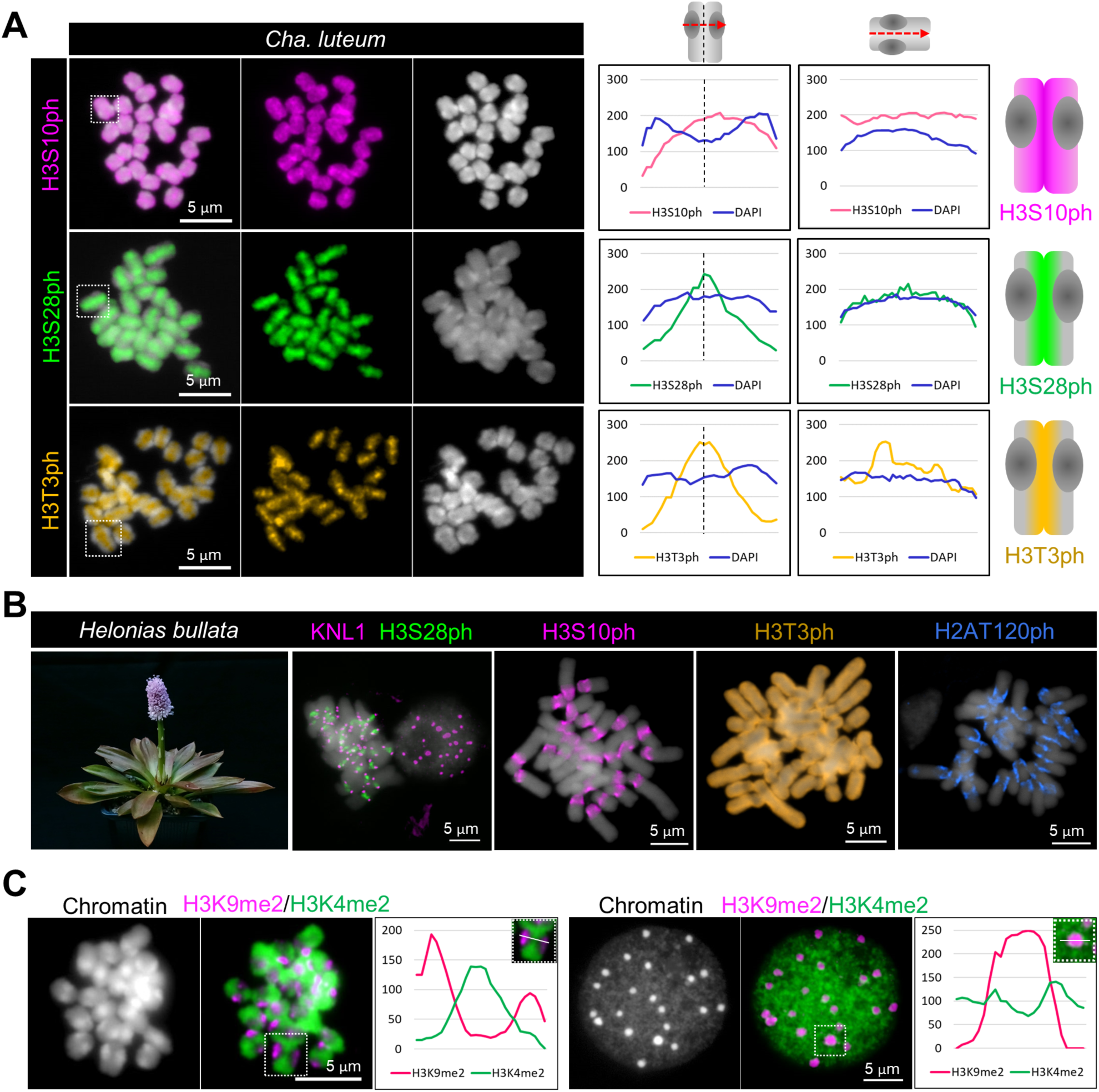
Visualization of cell cycle-dependent and eu- and heterochromatin-specific post-translational histone modifications. (A) Mitotic metaphase chromosomes of *Cha. luteum* after immunostaining with antibodies recognizing histone H3S10ph (magenta), H3S28ph (green) and H3T3ph (orange). The line scan plot profiles show the signal intensities of histone marks and DAPI measured in the framed chromosomes (squares). Signal distribution along single chromosomes is depicted as schemata next to the profiles. (B) Mitotic metaphase chromosomes of monocentric *Helonias bullata* after immunostaining with antibodies recognizing a combination of histone H3S28ph (green) and KNL1 (magenta), H3S10ph (magenta), H3T3ph (orange), and H2AT120ph (blue). (C) The immunolabelling patterns of H3K9me2 (magenta) and H3K4me2 (green) on metaphase chromosomes and interphase nuclei of *Cha. luteum* show the large-scale hetero- and euchromatin organization. (A–C) Chromosomes were counterstained with DAPI.

To determine whether the holocentromere-like histone phosphorylation patterns evolved before the formation of the tribe Chionographideae, we examined the chromosomal distribution of the phosphorylated histone variants in the sister tribe Heloniadeae (Figure 1A). Three species, *Helonias bullata*, *Heloniopsis umbellata*, and *Ypsilandra thibetica*, were selected as representatives of the corresponding genera. First, we confirmed the monocentricity of these three species by monitoring distribution of the conserved outer kinetochore protein KNL1 ^41^ (Figures 6B and S8). In these species, all analyzed histone H3 phosphorylation patterns were consistent with monocentricity and H2AT120ph signals were detectable at centromeric regions (Figures 6B and S8).

To probe the large-scale organization of eu- and heterochromatin in *Cha. luteum*, we assayed the evolutionarily conserved histone marks histone H3K4me2 and H3K9me2. The euchromatin mark H3K4me2 showed uniform signals in metaphase chromosomes, except at centromeres, where the heterochromatin histone mark H3K9me2 was enriched (Figure 6C). Similar labeling patterns were observed in interphase nuclei with H3K9me2 and H3K4me2 enriched in chromocenters and euchromatin, respectively (Figure 6C).

Thus, although *Cha. luteum* possesses a typical monocentric distribution of eu- and heterochromatin, it exhibits a cell cycle-dependent histone phosphorylation pattern remarkably similar to that of holocentric species, including *Chi. japonica*. This chromosome-wide phosphorylation pattern, unique to Chionographideae species and distinguishing them from Heloniadeae species with typical monocentromeres, likely evolved after the divergence of the two tribes.

## Discussion

### Unraveling centromere evolution in Chionographideae

Our analysis of atypical centromeres provides insights into the evolutionary transition between monocentricity and holocentricity. The constriction-free, macro-monocentromeres of the monocentric *Cha. luteum*, which is phylogenetically related to the holocentric *Chi. japonica*, shows both mono- and holocentric characteristics, which means that it is of unique value for investigating centromere evolution. We propose a model explaining the processes driving the divergence of centromere architectures in Chionographideae (Figure 7). In short, the divergent evolution of both atypical centromeres is a result of kinetochore gene mutations, alterations of mitotic histone phosphorylation patterns, and amplification of centromeric satellite DNA. In *Chi. japonica,* chromosome-wide spreading of centromere units resulted in the formation of a holocentromere. In *Cha. luteum,* local centromere expansion formed a macro-monocentromere.

**Figure 7.**
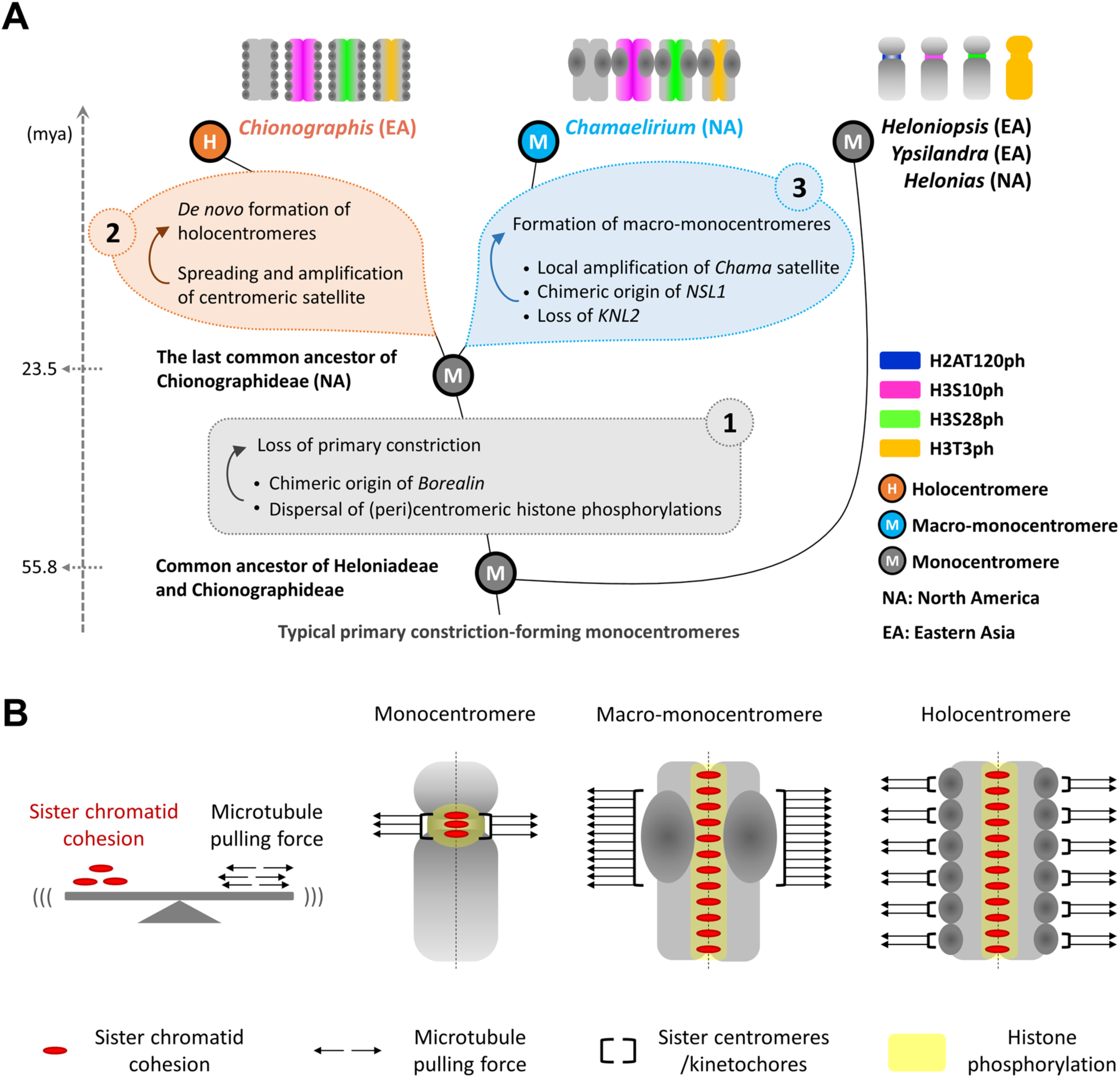
Model explaining the divergent evolution of holo- and macro-monocentromeres in Chionographideae. (A) The sister tribes Heloniadeae and Chionographideae, which diverged ∼55 million years ago (mya), exhibit intercontinental disjunctions between eastern Asia (EA) and eastern North America (NA). *Chionographis* and *Chamaelirium* possess primary constriction-free chromosomes with holocentromeres and macro-monocentromeres, respectively. In contrast, the monocentromeres of Heloniadeae species possess a primary constriction, suggesting that the loss of this chromosome constriction occurred before the divergence of holocentric *Chionographis* and monocentric *Chamaelirium* ∼23.5 mya. (1) The mutation of Borealin likely represents a molecular trigger leading to the chromosome-wide distribution of histone phosphorylation and driving the evolution of primary constriction-free chromosomes in Chionographideae. (2) Broad-scale genome synteny between *Chi. japonica* and *Cha. luteum* suggests a *de novo* origin of the holocentromeres in *Chi. japonica.* The evolution of both centromere-types was influenced by an amplification of centromeric repeats. In *Chi. japonica*, beside centromere-repeat amplification, chromosome-wide spread of centromere units resulted in the formation of a holocentromere. (3) In *Cha. luteum*, local centromere expansion resulted in macro-monocentromeres. However, the relationship between alterations in *KNL2* and *NSL1* and the local centromere expansion remains unclear. (B) Accurate chromosome segregation relies on a balance between sister chromatid cohesion and microtubule-generated pulling forces. As an adaptation required for proper chromosome segregation, the expansion of centromeric regions in both *Chamaelirium* and *Chionographis* probably increased microtubule attachment and pulling forces to counteract the increased sister chromatid cohesion caused by the chromosome-wide distribution of histone H3 phosphorylation. Macro-monocentromeres and holocentromeres represent two distinct outcomes of divergent evolution, each reflecting different adaptations to counteract the increased sister chromatid cohesion.

To elaborate, the sister tribes Heloniadeae and Chionographideae, which diverged ∼55 mya, exhibit intercontinental disjunctions between eastern Asia and eastern North America ^29^. Monocentric species exist in both tribes, implying a monocentric nature of their common ancestors. All *Heloniadeae* species examined possess a primary centromere constriction, suggesting that the loss of this defining chromosome structure occurred after the divergence of the two tribes, but before the evolution of the genera *Chamaelirium* and *Chionographis* at ∼23.5 mya (Figure 7A). While Heloniadeae species display cell cycle-dependent histone phosphorylation patterns typical for monocentricity, *Cha. luteum* and *Chi. japonica*, despite their different centromere structures, share similar distributions of histone H3S10ph, H3S28ph, and H3T3ph, which are typical for holocentricity, coupled with the absence of H2AT120ph. These findings suggest that alterations in (peri)centromeric histone phosphorylation sites also arose after the divergence of the Heloniadeae and Chionographideae lineages.

In addition to the similar histone phosphorylation patterns, the two species harbor the same mutation in the *Borealin* gene (Figure 7A-1). The Borealin protein – a component of the chromosomal passenger complex (CPC) – plays a crucial role in regulating histone phosphorylation ^44^. Hence, this change in the *Borealin* gene potentially represents a molecular trigger initiating chromosome-wide distribution of histone phosphorylation and driving the evolution of primary constriction-free chromosomes within Chionographideae.

The remarkably high proportion (∼15%) of centromeric satellite repeats in the genomes of both species suggests that the evolution of both centromere types was influenced by the amplification of centromeric repeats. The large-scale genome synteny between holocentric *Chi. japonica* and monocentric *Cha. luteum* suggests a *de novo* origin of holocentromeres in *Chi. japonica* (Figure 7A-2). However, the mechanisms underlying the chromosome-wide distribution of *de novo*-generated centromere units in holocentric *Chi. japonica* remain unknown ^3^. Local centromere expansion in *Cha. luteum* led to the formation of macro-monocentromeres, whose centromere proportion resembled that of holocentric *Chi. japonica* (Figure 7A-3). Given the chromosome-wide synteny and identical chromosome number in both Chionographideae species, multiple fusion events of monocentric chromosomes, as proposed for the holocentromeres in *Luzula* ^45^, is likely not the mechanism behind holocentromere formation in *Chionographis*.

Remarkably, the loss of the *KNL2* gene and the formation of chimeric *NSL1* occurred only in *Cha. luteum*, while both genes remain intact in *Chi. japonica*. Thus, the kinetochore composition of the primary constriction-free *Cha. luteum* macro-monocentromere appears to diverge more significantly from that of a typical monocentromere than does the holocentromere of *Chi. japonica*. This implies that both centromere types have evolved in parallel after diverging from their last common monocentric ancestor, rather than following a linear transition from monocentric *Chamaelirium* to holocentric *Chionographis*. Nevertheless, the relationship between alterations in KNL2 and NSL1 and the local expansion of centromeres in *Cha. luteum* remains unknown. It remains unclear whether centromere type changed gradually over time, as proposed for the stick insect ^46^, or resulted from a single event.

The centromere repeat monomers in both centromere types share a conserved 9-base pair sequence (TTCGTACGA). This sequence might facilitate the formation of non-B DNA hairpins, a DNA structure known to cause replication fork collapse and subsequent DNA double-strand breaks (DSB) ^47^. High DSB rates promote non-allelic (ectopic) recombination repair, which can reposition repeats at new genomic locations even over large distances ^48^. Non-B-form DNA-enriched centromeres may represent an ancient form of centromere specification, potentially through interaction with DNA-binding proteins that promote CENH3 loading ^49^. It remains unknown whether sequence-driven CENH3 loading occurs in Chionographideae.

### *Cha. luteum* exploits a KNL2-independent mechanism of CENH3 loading and centromere maintenance

Although the mechanism of CENH3 loading in plants is not yet fully understood, it likely depends on KNL2 ^50,51^. Thus, the absence of KNL2 in *Cha. luteum* might impair CENH3 loading at the centromeres. In *A. thaliana*, which has two KNL2 variants, αKNL2 mutants exhibited reduced levels of CENH3 at centromeres and showed mitotic and meiotic defects but were viable, whereas βKNL2 mutants exhibited complete lethality at the seedling stage ^50,51^. The importance of KNL2 for CENH3 loading onto centromeres has also been demonstrated in animals ^52–55^, suggesting that the role of KNL2 in CENH3 loading is evolutionarily conserved. Melanthiaceae species, like other monocots except grasses ^51^, possess only a single *KNL2* gene, which encodes a protein structurally more similar to αKNL2 of *A. thaliana*. Despite the loss of the *KNL2* gene, we observed no mitotic defects in *Cha. luteum,* suggesting that this species exploits an alternative, KNL2-independent mechanism for CENH3 deposition that specifically targets the centromeres. Two holocentric plant species of the genus *Cuscuta*, *C. europaea* and *C. epithymum*, have also been documented to have lost *KNL2* ^19^. In this study, the loss of both *αKNL2* and *βKNL2* genes was neither lethal nor led to mitotic defects and the authors hypothesized that CENH3 is no longer a centromeric protein, and the two *Cuscuta* species evolved a CENH3-independent mechanism for attachment to mitotic spindles to chromosomes ^19,56^. In *Cha. luteum*, this is likely not the case, as chromosomes bind to mitotic spindle microtubules exclusively at CENH3-containing domains, suggesting that CENH3 retains its role as a key centromere protein.

NASP, a general histone H3 chaperone present in both Chionographideae species, is the only protein currently known to participate in CENH3 deposition in plants ^57–59^. Although NASP, like KNL2, has been shown to bind to CENH3, the roles of the two proteins differ. While KNL2 ensures CENH3 loading onto centromeres via an interaction with centromeric nucleosomes, NASP does not directly participate in CENH3 deposition. Instead, NASP binds non-nucleosomal CENH3 and escorts it to chromatin assembly factor(s) ^57,58^. Thus, it remains unclear which alternative mechanism is responsible for CENH3 loading in *Cha. luteum*.

Interestingly, all three species lacking *KNL2* have unusual centromeres. Centromeric chromatin in *Cha. luteum* expanded enormously at loci containing highly amplified satellite DNA, comprising up to 15 Mb DNA per centromere. CENH3-containing heterochromatin domains in *Cuscuta europaea* are present at one to three sites per chromosome and are closely associated with the satellite DNA family CUS-TR24, which spans in total 181.2 Mb, corresponding to an average of 25 Mb per chromosome ^19^. In contrast, no detectable CENH3 is present on chromosomes in *C. epithymum*, which contains a low amount of satellite DNA ^19^. This shows that the loss of KNL2 has different consequences in different species, possibly dependent on the association of CENH3 with satellite DNA. Conversely, the observation of active *KNL2* in the holocentric *Chi. japonica* suggests that the absence of this gene is not a prerequisite for the evolution of holocentromeres across species.

### The possible impact of the chimeric Borealin and NSL1

Chimeric genes are important players in the evolution of genetic novelty ^60^. Three ancient domain fusions between kinetochore proteins were predicted to occur in the last common ancestor of eukaryotes ^61^. The chimeric origin of NSL1 and Borealin in Chionographideae is unprecedented among kinetochore protein genes.

Borealin is a component of the chromosome passenger complex (CPC), which is a key regulator of mitotic events ^62^. The Borealin N-terminus is required for the interaction of CPC with nucleosomes ^44^. In *Cha. luteum* and *Chi. japonica* the N-terminus of Borealin has been replaced by a 50 amino acid-long fragment of RPS17. CPC binding to nucleosomes is an upstream requirement for Haspin (phosphorylates H3T3) and Bub1 (phosphorylates H2AT120) activities, and for Haspin/Bub1-mediated CPC enrichment at centromeres. Further, phosphorylated H3T3 is required for centromeric recruitment of the CPC component Aurora B, which phosphorylates H3S10 and H3S28 ^42,63^. Possibly, a mutation of Borealin that resulted in a chimeric protein led to the observed chromosome-wide distribution of H3S10ph and H3S28ph and the absence of H2AT120ph. Consequently, misregulation of this pathway could disrupt pericentromeric modifications, further affecting chromatid cohesion and chromosome segregation.

NSL1, as a component of MIS12c, is one of the key structural kinetochore proteins. It is essential for interaction with the other two complexes of the outer kinetochore, NDC80c and KNL1c ^39^. NSL1 knockout mutants were lethal in Drosophila ^64^. Lack of KNL1 and NDC80 immunosignals indicate that these proteins are absent or else expressed only at very low levels, or that the fusion of NSL1 with the N-terminal region of BORCS6 disrupts the formation of the outer kinetochore KMN complex in centromeres of *Cha. luteum*.

BORCS6 belongs to the BLOC-one-related complex (BORC) ^65^. In animal cells, BORC controls lysosomal and synaptic vesicle transport and positioning by recruiting ARL8, which either directly interacts with kinesin-3 or indirectly associates with kinesin-1 through cargo adaptors, coupling lysosomes with kinesin motors ^65^. Although plant cells do not contain lysosomes, BORC ^66^, ARL8 ^67^ and kinesin ^68^ protein homologs are present, and thus BORC might be involved in organelle or vesicle transport as well. It is tempting to speculate that the fusion of BORCS6 N-terminus with the C-terminus of NSL1 could recruit BORC to the kinetochore. If true, chimeric BORC could bring additional function(s) to the kinetochore in *Cha. luteum*.

Since only 29 structural and regulatory kinetochore genes were considered in our comparative study, we cannot exclude that additional changes in kinetochore complex were involved in the evolution of holo- and macro-monocentromeres.

### Centromere diversity: different paths to a common functional adaptation

The existence of constriction-free centromeres in *Cha. luteum* suggests that – although monocentromeres are often distinguished by a primary constriction – this structure is not always necessary for centromere function. The folding of centromeric chromatin ensures that CENH3-containing nucleosomes are oriented poleward, facilitating accurate chromosome segregation. The absence of a primary constriction in this species may be explained by the holocentromere-typical distribution of histone H3 phosphorylation marks along the entire length of the chromosomes and the unusually large centromere size.

Accurate chromosome segregation relies on a delicate balance between sister chromatid cohesion and microtubule-generated pulling forces ^69^ (Figure 7B). Consequently, as an adaptation required for proper chromosome segregation, the expansion of centromeric regions in both *Chamaelirium* and *Chionographis* probably increased microtubule attachment and pulling forces to offset the increased sister chromatid cohesion brought on by the chromosome-wide distribution of pericentromeric histone H3 phosphorylation. Macro-monocentromeres and holocentromeres represent distinct outcomes of divergent evolution and different adaptation to counteract the increased sister chromatid cohesion. These divergent solutions highlight the plasticity of centromere evolution in response to selective pressure imposed by changes in chromatid cohesion dynamics driven by mutation in a single gene (chimeric Borealin). Moreover, these findings suggest a complex interplay between centromere size, number and distribution of centromere units (mono versus holo), and chromatid cohesion in shaping chromosome morphology. Overall, the two Chionographideae species offer valuable insights into centromere type evolution, highlighting the importance of incorporating greater phylogenetic diversity in model organisms used to study fundamental chromosomal features.

## Materials and Methods

### Plant materials

*Chamaelirium luteum* (L.) A. Gray plants used in this study were provided by the Deutsche Homöopathie-Union (DHU), Germany, *Chionographis japonica* (Willd.) Maxim. plants were obtained from commercial nurseries in Japan, and the *Helonias bullata* L., *Heloniopsis umbellata* Baker, *Heloniopsis orientalis* var. *breviscapa*, and *Ypsilandra thibetica* Franch. species were purchased from British nurseries in the UK. The plants were grown at IPK Gatersleben (Germany) in a greenhouse: 16 h light (from 6 AM to 10 PM), day temperature 16 °C, night temperature 12 °C. Seeds of *Sorghum bicolor* BTx623 and *Secale cereale* L. inbred line Lo7 were obtained from the IPK GeneBank (Gatersleben, Germany) and were germinated on wet filter papers to harvest roots and young leaves for experiments.

### Genome size measurement

To isolate nuclei, approximately 0.5 cm^2^ of fresh leaf tissue of *Cha. luteum* was chopped together with equivalent amounts of leaf tissue of either of the two internal reference standards *Glycine max* (L.) Merr. convar. max var. max, cultivar ‘Cina 5202’ (Gatersleben genebank accession number: SOJA 392; 2.21 pg/2C) or *Raphanus sativus* L. convar. sativus, cultivar ‘Voran’ (Gatersleben genebank accession number: RA 34; 1.11 pg/2C), in a petri dish using the reagent kit ‘CyStain PI Absolute P’ (Sysmex-Partec) following the manufacturer’s instructions. The resulting nuclei suspension was filtered through a 50-μm CellTrics filter (Sysmex-Partec) and measured on a CyFlow Space flow cytometer (Sysmex-Partec, Germany). At least six independent measurements were performed for *Cha. luteum*. The absolute DNA content (pg/2C) was calculated based on the values of the G1 peak means and the corresponding genome size (Mbp/1C), according to ^70^.

### Short-read sequencing of DNA and RNA

Genomic DNA of *Cha. luteum* was extracted from leaf tissue using the DNeasy Plant Mini kit (Qiagen, Germany). Low-pass paired-end (2×150 bp) genome sequencing was performed using DNBSEQ system performed by BGI (China). Total RNAs from leaf, root, and fruit tissues of *Cha. luteum* were isolated using the Spectrum^TM^ Plant total RNA kit (Sigma, USA, cat. no. STRN50). Library preparation (Illumina Stranded mRNA Prep Ligation Kit) and sequencing at IPK Gatersleben or Novogene (UK) (paired-end, 2 × 151 cycles, Illumina NovaSeq6000 system) involved standard protocols from the manufacturer (Illumina Inc., USA).

### Isolation of HMW DNA, HiFi library preparation, and sequencing

For long-read PacBio sequencing, high-molecular weight (HMW) DNA of *Cha. luteum* was isolated from leaves of a single plant using the NucleoBond HMW DNA kit (Macherey Nagel, Germany). Quality was assessed using the FEMTO Pulse system (Agilent Technologies Inc, CA, USA). Quantification involved the Qubit device and the dsDNA High Sensitivity assay kit (Thermo Fisher Scientific, MA, USA). A HiFi library was prepared from 15 µg HMW DNA using the “SMRTbell prep Kit 3.0” according to the manufacturer protocol (Pacific Biosciences of California Inc., CA, USA). The initial DNA fragmentation was performed using the Megaruptor 3 device (Shear speed: 29; Diagenode, Belgium). Finally, HiFi libraries were size-selected (narrow-size range: approximately 20 kb) using the SageELF system with a 0.75% Agarose Gel Cassette as described by the manufacturer (Sage Science Inc., MA, USA). Sequencing (HiFi CCS) was performed using the Pacific Biosciences Revio device (24 h movie time, 155 pM loading concentration, 2 h pre-extension time, diffusion loading, 100 min loading time, 22 kb mean insert length according to SMRT link raw data report, 55 Gb HiFi CCS yield) following standard manufacturer’s protocols (Pacific Biosciences of California Inc., Menlo Park, CA, USA) at IPK Gatersleben.

### Chromosome conformation capture (Hi-C) sequencing

Hi-C sequencing libraries were generated from flowers of *Cha. luteum* as described previously ^71^ using *Dpn*II enzyme, and were sequenced (paired-end, 2×111 cycles) using the NovaSeq6000 device (Illumina Inc., USA) at IPK Gatersleben. A total of ∼102 Gb paired-end reads were generated. After filtering, ∼87 Gb Hi-C read pairs were used for scaffolding.

### Genome assembly and Hi-C scaffolding

A total of ∼53 Gb of PacBio HiFi reads (∼59.5× coverage) of *Cha. luteum* were assembled into contigs using hifiasm (v0.19.3-r572; default) ^72^. Contig statistics were calculated with Quast (v2.3) ^73^ and gene content completeness was evaluated with Benchmarking Universal Single-Copy Orthologs (BUSCO) (v4.1.2; dataset: liliopsida_odb10, E-value=0.001) ^74^. An initial 1.56-Gb primary assembly of *Cha. luteum* genome was constructed with a BUSCO completeness of 95.6% (Table S1). The unexpectedly large assembly size and the high score of duplicated BUSCOs (68.0%) suggested the presence of notable heterozygosity in the primary contigs. To join the residual heterozygous contains, purge_dups (v 1.2.5; default) ^75^ was used to identify and remove duplicated sequence segments in the primary assembly, resulting in duplicated BUSCOs reduced to 10.6% (Table S1). The Arima Genomics mapping pipeline (https://github.com/ArimaGenomics/mapping_pipeline) was used to process the Hi-C data, including read mapping to the contigs, read filtering, read pairing, and PCR duplicate removal, and scaffolding was performed using YaHS (v1.2a.2; -e GATC --no-contig-ec) ^76^. Hi-C contact maps and manual curation were accomplished by the bash scripts provided (https://github.com/c-zhou/yahs) and visualized using Juicebox (https://github.com/aidenlab/Juicebox).

### Gene-based synteny analysis

The genome of *Chi. japonica* was re-annotated by mapping the clean RNA-seq data (EMBL ENA <u>PRJEB58123</u>) generated from our previous study ^31^. For genome-directed transcriptome assembly, the RNA-seq datasets generated in this study from fruits (∼7 Gb), leaves (∼13 Gb), and roots (∼13 Gb) of *Cha. luteum* (EMBL ENA PRJEB82608), were aligned to the assembled scaffolds using HISAT2 (v2.2.1, default parameters) ^77^, and then processed to produce gene feature annotation with StringTie (v2.1.1, default parameters) ^78^. Two sets of non-redundant transcripts from *Chi. japonica* and *Cha. luteum* were generated using gffread (v0.12.6) ^79^. TransDecoder (v5.5.0) ^80^ was used to annotate coding regions in transcripts. BRAKER3 pipeline was used to improve the annotation accuracy of protein-coding genes ^81^.

To find the links of conserved single-copy proteins between the *Chi. japonica* and *Cha. luteum* genomes, a <u>Python script</u> was developed as follows. First, the translated protein sequences of *Chi. japonica* were aligned to those of *Cha. luteum* via blastp (v2.5.0, default). Second, alignments with an identity of < 90% and an alignment length of < 150 aa were treated as noisy alignments and were filtered out. Third, only when a sequence showed similarity to a single gene in the other genome were considered as conserved single-copy proteins. Finally, the genomic positions of the syntenic single-copy proteins were extracted from the gff3 files to create the links between the two genomes. NGenomeSyn (v1.41) ^82^ was used to visualize the genome synteny.

### Transcriptome-based identification of kinetochore protein genes

The clean RNA-seq datasets from various tissues of *Cha. luteum* were *de novo* assembled using Trinity 2.4.0 ^80,83^ with default parameters. In addition, the genome-directed assembled transcriptome of *Cha. luteum* as described above was also applied for downstream analyses. Putative coding regions were first identified by TransDecoder (v5.5.0) ^80^ with a minimum protein length of 100 as a threshold.

Trinity-made transcriptome assembly of *Chi. japonica* was obtained from our previous study ^31^. The RNA-seq data and genome assembly used in the study were applied here for genome-directed transcriptome assembly using HISAT2 (v2.2.1, default) ^77^ followed by StringTie (v2.1.1, default) ^78^, as described above. *De novo* transcriptome assembly in *H. orientalis* was constructed using RNA-seq data downloaded from Sequence Read Archive (https://www.ncbi.nlm.nih.gov/sra; SRR28160651 and SRR28160654).

Protein sequence databases for these species were constructed by translating predicted open reading frames from *de novo* and genome-directed transcriptome assemblies generated using Trinity (all three species) and StringTie (*Chi. japonica* and *Cha. luteum),* respectively. The analysis was done using two types of sequence data; gene annotation models predicted in genome assemblies based on RNA-seq data and transcriptome assemblies.

Kinetochore protein sequences were first identified in *P. dactylifera* using BLASTp searches in the GenBank protein database, guided by sequences from previous studies ^19,37,84,85^. These identified sequences were then used to search for homologs in the three Melanthiaceae species. The sequences of kinetochore proteins translated from *de novo* and genome-derived transcriptome assemblies matched well. Since the genome-derived assemblies were coupled with detailed information about the gene structures, we used these for further analyses. In a few cases, the gene structures had to be manually corrected by reassembling the RNA-seq reads assigned to specific loci and using the programs est2genome ^86^ and genewise ^87^ for the gene annotations.

### Phylogenetic analysis

The CENH3 protein sequences of *Cha. luteum* and the other species derived from the NCBI GenBank (Table S6) were aligned using the ClustalW algorithm implanted in MEGA X by default setting ^88,89^. The maximum-likelihood tree was constructed via IQ-Tree web server (http://iqtree.cibiv.univie.ac.at/) ^90^ and visualized using Interactive Tree Of Life (iTOL, http://itol.embl.de/) ^91,92^.

### Antibody production

The synthesized peptides of *Cha. luteum*CENH3 (ClCENH3: MAPTKKTKKTTENINNRPAL-C) were used for the immunization of rabbits to generate polyclonal antibodies. The peptide synthesis, immunization, and antibody purification were performed by LifeTein (www.lifetein.com, USA).

### Indirect immunostaining

Mitotic chromosomes and interphase nuclei were prepared from root meristems. Roots were pretreated in ice-cold water overnight and fixed in 4% paraformaldehyde in Tris buffer (10 mM Tris, 10 mM EDTA, 100 mM NaCl, 0.1% Triton X-100, pH7.5) for 5 min on ice under vacuum treatment, followed by another 25–30 min solely on ice. Root meristems were then chopped in lysis buffer LB01 (15 mM Tris, 2 mM Na_2_EDTA, 0.5 mM spermine, 80 mM KCl, 20 mM NaCl, 15 mM β-mercaptoethanol, and 0.1 % (v/v) Triton X-100) ^93^, the cell suspension was filtered through a 50-μm CellTrics filter (Sysmex-Partec) and subsequently centrifuged onto slides using a Cytospin3 (Shandon, Germany) at 700 rpm (×55.32 g) for 5 min. The chromosome spreads were blocked in 3% BSA in 1× phosphate-buffered saline (PBS) at room temperature (RT) for 1 h and incubated with primary antibodies in 1% BSA/1× PBS at 4°C overnight. After three washes in 1× PBS at RT for 5 min each, secondary antibodies in 1% BSA/1× PBS were applied, followed by an incubation at 37°C for 1 h. After three washes, the slides were dehydrated in 70-90-100% ethanol series for 3 min each and counterstained with 10 µg/ml 4’,6-diamidino-2-phenylindoline (DAPI) in Vectashield antifade medium (Vector Laboratories, USA). For immunodetection of microtubules, root pretreatment with ice-cold water was omitted, and the Tris buffer and 1× PBS mentioned above were substituted by 1× MTSB buffer (50 mM PIPES, 5 mM MgSO_4_, and 5 mM EGTA, pH 7.2).

The primary antibodies used in this study included customized rabbit anti-*Cha. luteum* CENH3 (dilution 1:500), rabbit anti-*Chi. japonica* MIS12 (dilution 1:100) ^31^, rabbit anti-*Chi. japonica* NDC80 (dilution 1:100) ^31^, and rabbit anti-*Cuscuta europeae* KNL1 (dilution 1:500 or 1:1000) ^19^, as well as the commercially available mouse anti-alpha-tubulin (Sigma-Aldrich, USA, cat. no. T9026-2, dilution 1:300), rabbit anti-histone H3K4me2 (abcam, UK, cat. no. ab7766, dilution 1:300), mouse anti-histone H3K9me2 (abcam, UK, cat. no. ab1220, dilution 1:300), mouse anti-histone H3S10ph (abcam, UK, cat. no. ab14955, dilution 1:1000), rat anti-histone H3S28ph (Sigma-Aldrich, USA, cat. no. H9908, dilution 1:1000), rabbit anti-H3T3ph (Sigma-Aldrich, USA, cat. no. 07-424, dilution 1:1000), and rabbit anti-H2AT120ph (Active Motif, USA, cat. no. 61196, dilution 1:500).

The anti-rabbit rhodamine (Jackson ImmunoResearch, USA, cat. no. 111-295-144, dilution 1:300), anti-rabbit Alexa488 (Jackson ImmunoResearch, USA, cat. no. 711-545-152, dilution 1:300), anti-mouse Alexa488 (Jackson ImmunoResearch, USA, cat. no. 715-546-151, dilution 1:300), and anti-rat Alexa488 (Jackson ImmunoResearch, USA, cat. no. 112-545-167, dilution 1:300) were used as secondary antibodies.

### Repeatome analysis

The low-coverage genome skimming dataset of *Cha. luteum* was generated in this study, and those of *Chi. japonica* (ERR10639507, EMBL ENA, https://www.ebi.ac.uk/ena/) ^31^ and *H. umbellata* (SRR15208642, NCBI SRA, https://www.ncbi.nlm.nih.gov/sra/) ^94^ were publicly available. Genomic PE reads were assessed by FastQC ^95^ implanted in the RepeatExplorer pipeline (https://repeatexplorer-elixir.cerit-sc.cz/galaxy/) and filtered by quality with 95% of bases equal to or above the cut-off value of 10. Qualified PE reads of *Cha. luteum* equivalent to 0.5× genome coverage were applied to analyze repetitive elements by a graph-based clustering method using RepeatExplorer ^96–98^. The automatic annotation of repeat clusters was manually inspected and revised if necessary, followed by a recalculation of the genome proportion of each repeat type. The comparative clustering analysis was performed based on one million PE reads from each of the three species.

### Chromatin immunoprecipitation (ChIP) sequencing

The ChIP experiment was performed with minor modifications as described by ^31^. 0.65 g of *Cha. luteum* flower and 1.0 g of *Secale cereale* (inbred line Lo7) leaf tissue were ground with liquid nitrogen and homogenized separately in 10 ml nuclei isolation buffer (1 M sucrose, 5 mM KCl, 5 mM MgCl_2_, 60 mM HEPES pH 8.0, 5 mM EDTA, 0.6% Triton X-100, 0.4 mM PMSF, 1 µM pepstatin A, cOmplete protease inhibitor cocktail (Roche)). Nuclei fixation was performed in 1% PFA/nuclei isolation buffer at RT, 12 rpm for 10 min and terminated by adding glycine to a final concentration of 130 mM. The nuclei suspension was filtrated through Miracloth (Millipore) twice and a 50-µm CellTrics filter (Sysmex) once and centrifuged at 4°C, 3,000 ×g for 10 min. The nuclei pellet was resuspended in 1 ml extraction buffer (0.25 M sucrose, 10 mM Tris-HCl pH 8.0, 10 mM MgCl_2_, 1% Triton X-100, 1 mM EDTA, 5 mM β-mercaptoethanol, 0.1 mM PMSF, 1 µM pepstatin A, cOmplete protease inhibitor cocktail), followed by centrifugation at 4°C, 12,000 ×g for 10 min. After removing the supernatant, nuclei were resuspended in 150 µl of nuclei lysis buffer (20 mM Tris-HCl pH 8.0, 10 mM EDTA, 1% SDS, 0.1 mM PMSF, 1 µM pepstatin A, cOmplete protease inhibitor cocktail). Chromatins were sonicated for 14 cycles of 30 s ON, 30 s OFF at high power in a Bioruptor (Diagenode), followed by an addition of 100 µl ChIP dilution buffer (16.7 mM Tris-HCl pH 8.0, 167 mM NaCl, 1.1% Triton X-100, 1 mM EDTA, cOmplete protease inhibitor cocktail), and continued sonication to a total of 31 cycles under the same setting. The sonicated samples were diluted 10 times with ChIP dilution buffer, centrifuged at 4°C, 13,000 ×g for 5 min, and the supernatant of each sample was transferred to new tubes. To dilute the high proportion of the putative *Cha. luteum* centromeric repeat, sonicated chromatin *of Cha. luteum* and *S. cereale* were mixed in a 1:3 ratio. The mixed chromatins were incubated with the ClCENH3 antibody (10 mg/ml) to a final 1:500 dilution at 4°C by shaking at 14 rpm for 12 h. Dynabeads^TM^ Protein A (Invitrogen) in ChIP dilution buffer, corresponding to 0.1× volume of the chromatin solution, was added to the antibody-prebound chromatins and incubated at 4°C by shaking at 14 rpm for 1.5 h. The collected beads were then washed twice in low salt buffer (150 mM NaCl, 0.1% SDS, 1% Triton X-100, 2 mM EDTA, 20 mM Tris-HCl pH 8.0), followed by three washes in high salt buffer (500 mM NaCl, 0.1% SDS, 1% Triton X-100, 2 mM EDTA, 20 mM Tris-HCl pH 8.0), and another two washes in TE buffer at 4°C by shaking at 14 rpm for 5 min. The bead-bound chromatin was purified by using iPure kit v2 (Diagenode) following the manual and quantified using Qubit^TM^ dsDNA HS Assay kit (Invitrogen). ChIPseq libraries were prepared by NEBNEXT^®^ Ultra^TM^ II DNA Library Prep Kit for Illumina (New England Biolabs) and sequenced using NovaSeq 6000 system (Illumina) by Novogene (UK) in the paired-end run (2×150 bp).

### ChIP-seq data analysis

To evaluate the enrichment of repeats associated with CENH3-containing nucleosomes, single-end reads of CENH3-ChIP-seq and input-seq were quality filtered using the tool ‘’Processing of FASTQ reads’’ (Galaxy Version 1.0.0.3), implanted in the Galaxy-based RepeatExplorer portal (https://repeatexplorer-elixir.cerit-sc.cz/galaxy/). ChIP-Seq Mapper (Galaxy version 1.1.1.4) (Neumann et al., 2012) was used to map the ChIP- and input reads on RepeatExplorer-derived contig sequences of repeat clusters. To analyze the size and position of *Cha. luteum* centromeres, the paired-end reads of ChIP- and input-seq were quality-filtered by Trimmmomatic (Galaxy Version 0.39) ^99^ and the resulting reads were mapped to the *Cha. luteum* genome assembly using Bowtie2 (Galaxy Version 2.5.3) ^100^ with default parameters. The deeptools bamCompare function (Galaxy Version 3.5.4) ^101^ was used to generate the normalized ChIP-seq signal track of the average of read counts in ChIP over input in genome-wide 1 kb windows. Plots of chromosome regions with multiple tracks were plotted with pyGenomeTracks (Galaxy version 3.8) ^102^. To generate the sequence identity heatmap, the StainedGlass package was used ^103^.

### ATAC sequencing and data analysis

Fresh leaf of *Cha. luteum* were finely chopped using a razor blade in 1 ml of nuclei isolation buffer (0.25 M sucrose, 10 mM Tris-HCl pH 8.0, 10 mM MgCl_2_, 1% Trion X-100, 5 mM β-mercaptoethanol) supplemented with 1× Halt™ Protease Inhibitor Cocktail (Thermo Scientific), and the slurry was filtered through a 50-µm cell strainer. The resulting nuclei were washed twice and resuspended with the same nuclei isolation buffer. An aliquot was analyzed using a flow cytometer for quality control and quantification. Based on the quantification, a volume containing approximately 75,000 nuclei was aliquoted, and the nuclei pellet was collected by centrifugation.

The nuclei pellet was resuspended in a transposition reaction mix containing Tagment DNA Enzyme (TDE1, Illumina, 20034197), 0.4× PBS, 0.01% digitonin, and 0.1% Tween-20, and incubated at 37°C for 30 min. Transposition products were purified using the MinElute PCR Purification Kit (QIAGEN, 28004). Libraries were amplified with the NEBNext® High-Fidelity 2× PCR Master Mix (NEB, M0541), and further purified using VAHTS^TM^ DNA Clean Beads (Vazyme, N411). The final libraries were sequenced in paired-end mode (2× 151 cycles) on the Illumina NovaSeq 6000 (Illumina Inc., USA).

Sequencing reads were adapter-trimmed using fastp (v0.20.0) ^104^ and aligned to the reference genome with BWA-MEM (v0.7.17) ^105^. SAMtools (v1.16.1) ^106^ was used to remove duplicate and multi-mapping reads (-q 30), followed by peak calling (-q 0.01) with MACS (v3.0.0) ^107^. For visualization, BAM files containing uniquely mapped reads were converted to BigWig format using deepTools bamCoverage (v3.5.1) ^101^, with Reads Per Kilobase per Million mapped reads (RPKM) normalization.

### Preparation of fluorescence *in situ* hybridization (FISH) probes

The consensus sequences of satellite repeats reconstructed by TAREAN (TAndem REpeat ANalyzer) ^108^ were used to design fluorescence-modified oligonucleotides which were synthesized by Eurofins (Germany). The clones pAtT4 ^109^ was used as the probe to detect *Arabidopsis*-type telomeres. Plasmid DNA was labeled with ATTO550-dUTP using Fluorescent Nick Translation Labeling kits (Jena Bioscience, Germany).

The chromosome 2-specific oligo painting probes were designed based on the genome assembly of *Chi. japonica* ^31^ using the software Chorus2 ^110^. The predicted *Chionographis*-based oligos which matched to the genome of *Cha. luteum* were all included in the synthesized myTags Immortal Libraries (Daicel Arbor Biosciences, USA) to improve the probe transferability to *Cha. lutuem*. Oligo pools were labelled with fluorophores ATTO-594, Alexa 488, or Alexa 647 following the myTags Immortal Labeling Protocol (Daicel Arbor Biosciences, USA, https://arborbiosci.com/wp-content/uploads/2022/05/DaicelArborBio_myTags_Labeling_Protocol_v2-2.pdf).

### Chromosome preparation and FISH

Mitotic chromosome spreads were prepared from root meristems using a dropping method ^31^. Roots were pretreated in ice-cold water overnight, fixed in 3:1 (ethanol: glacial acetic acid) fixative at RT, overnight and kept in 70% ethanol at -20°C until use. Fixed roots were digested in an enzyme mixture (0.7 % cellulose Onozuka R10 (Duchefa Biochemie, The Netherlands, cat. no. C8001), 0.7 % Cellulase (Calbiochem, USA, cat. no. 219466), and 1.0 % pectolyase (Sigma, USA, cat. no. 45-P3026)) in citric buffer (0.01 M sodium citrate dihydrate and 0.01 M citric acid) at 37°C for 30–40 min. Cell suspension in the 3:1 fixative was dropped onto slides on a hot plate at 55°C, and slides were further fixed in 3:1 fixative for 1 min, air-dried, and kept at 4°C for later use.

To prepare meiotic chromosomes, inflorescences of *Chi. japonica* and *Cha. luteum* were fixed as described above for roots. Anthers were digested at 37°C for 70-80 min in an enzyme mixture (0.23 % cellulose Onozuka R10 (Duchefa Biochemie, The Netherlands, cat. no. C8001), 0.23 % Cellulase (Calbiochem, USA, cat. no. 219466), 0.33 % pectolyase (Sigma, USA, cat. no. 45-P3026), and 0.33 % cytohelicase (Sigma, USA, cat. no. C8247)). Meiotic spreads were prepared by a typical squash method ^111^. FISH mapping was performed as described in ^35^. For oligo-FISH, the hybridization mixture containing 10% dextran sulfate, 50% formamide, 2× SSC, and 500–1000 ng of each labeled oligo pool was used, and the hybridization at 37°C was extended to 36–48 h.

### Microscopy and image analysis

Widefield fluorescence images were captured using an epifluorescence microscope BX61 (Olympus) equipped with a CCD camera (Orca ER, Hamamatsu, Japan) and pseudo-colored by the Adobe Photoshop 6.0 software. To analyze the chromatin and centromere ultrastructures at the super-resolution level, we applied structured illumination microscopy (3D-SIM) using a 63x/1.40 Oil Plan-Apochromat objective of an Elyra 7 microscope system (Carl Zeiss GmbH, Germany). Image stacks were captured separately for each fluorochrome using 561, 488, and 405 nm laser lines for excitation and appropriate emission filters ^112^. Maximum intensity projections from image stacks were calculated using the ZENBlack software. Zoom-in sections were presented as single slices to indicate the subnuclear chromatin structures at the super-resolution level. 3D rendering to produce spatial animations was done based on 3D-SIM image stacks using the Imaris 9.7 (Bitplane, UK) software.

### Transmission electron microscopy (TEM)

For electron microscopy analysis, cuttings of root tips of 3 mm length were used for aldehyde fixation, dehydration and resin embedding as described (Kuo et al. 2023). Ultra-thin sectioning and TEM analysis were performed as explained previously ^113^

## Resource Availability

### Lead contact

Requests for further information and resources should be directed to and will be fulfilled by the lead contact, Andreas Houben

### Data and code availability

The whole-genome sequencing and RNA-seq datasets generated for this study can be found at EMBL-ENA under the project IDs PRJEB82607 and PRJEB82608, respectively. The datasets of ChIP-Seq (Project ID: PRJNA1201173, accession GSE285103) and ATAC-Seq (Project ID: PRJNA1201177, accession GSE285102) were deposited in the NCBI GEO database. The gene annotation, syntenic genes, and sequences of oligo-FISH painting probes and proteins are available in Zenodo (https://zenodo.org/records/15182433). The Python scripts for genome and synteny analyses are available in BitBuckket (https://bitbucket.org/ipk-csf/chamaelirium2chiographis/).

## Acknowledgements

We thank Oda Weiss, Ines Walde, Manuela Knauft, Susanne König, and Pascal Jaroschinsky (IPK) for technical help in DNA isolation and sequencing, Anne Fiebig (IPK) for sequence submission, Kirsten Hoffie and Marion Benecke (IPK) for technical support in sample preparation for transmission electron microscopy, and Takayoshi Ishii (Arid Land Research Center, Tottori University, Japan) for providing plant materials; Klaus Mayer (Helmholtz Center Munich, Munich, Germany), André Marques (MPI Breeding Research, Cologne, Germany) and Martin Mascher (IPK) for suggestion on genome assembly; Ingo Schubert (IPK) and Noriyuki Tanaka (Tokyo, Japan) for discussion; and Neysan Donnelly for editorial suggestions.

This work was supported by the Deutsche Forschungsgemeinschaft DFG grants No. HO1779/32-1 and HO1779/32-2 (to AH); the Taiwan Ministry of Science and Technology (MOST) grants MOST 106-2313-B-002-034-MY3 and MOST 108-2811-B-002-608) (to YTK); the Czech Science Foundation grant 25-15809S (to PN); the China Scholarship Council CSC202006850005 (to JC); the scholarships from National Council for Scientific and Technical Research (CONICET) and the National Agency for the Promotion of Science and Technology (FONCyT) project PICT 2020–1777 (to MAS).

## Author contributions

YTK performed the majority of the experiments, sequence and image analyses, including DNA and RNA isolation, immunostaining, FISH, ChIP-seq,; PN and JM performed kinetochore protein and genome sequence analyses; JC performed genome assembly, gene annotation, and synteny analysis; VS performed super-resolution microscopy and image analysis; KK performed immunostaining; JF conducted flow cytometry; MM performed electron microscopy; MAS performed FISH and data analysis; ZZ performed ATAC-seq and data analysis; AHim performed PacBio and Hi-C library preparation and sequencing, HH grew and provided plant materials; AH supervised the research project; YTK and AH wrote the manuscript with the input from all coauthors.

## Declaration of interests

The authors declare no competing interests.

## Supplemental Information

Document S1. Figures S1–S8 Document S2. Tables S1–S6

Video S1 Immunostaining of CENH3 (magenta) and spindle microtubules (green) in metaphase chromosomes of *Cha. luteum*.

## Supplemental Information Document S1: Figures S1–S8

**Figure S1.**
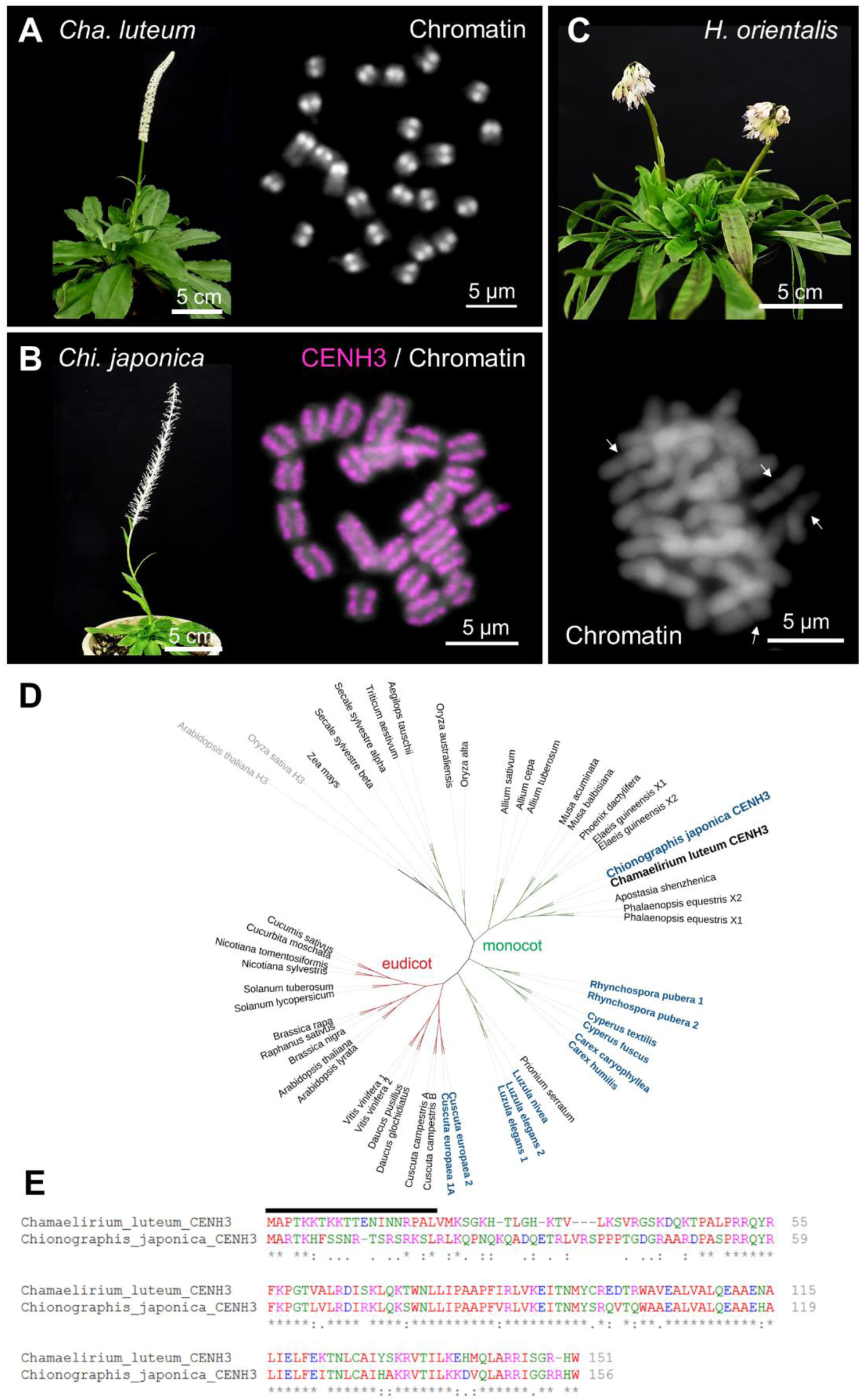
Diversity of chromosome and centromere structures in three Melanthiaceae species and identification of *Chamaelirium luteum* CENH3. (A) Flowering *Cha. luteum* features large heterochromatic monocentromeres that protrude at mitotic metaphase. The monocentromere-typical primary construction is missing in this species. (B) Flowering holocentric *Chionographis japonica* possesses chromosome-wide CENH3-immunosignals at the two peripheries of metaphase chromosomes (magenta). (C) Flowering *Heloniopsis orientalis* has monocentric chromosomes with a typical primary constriction (arrows). Chromosomes were counterstained with DAPI. (D) Phylogenetic tree of mono- and eudicot CENH3 proteins. CENH3 of monocentric *Cha. luteum* grouped with the CENH3 of holocentric *Chi. japonica.* Histone H3 of *Arabidopsis thaliana* and *Oryza sativa* was used as an outgroup. (E) Amino acid sequence alignment of CENH3s from *Cha. luteum* and *Chi. japonica*. The peptide sequence used for the generation of the *Cha. luteum*-specific anti-CENH3 antibody is indicated with a black line.

**Figure S2.**
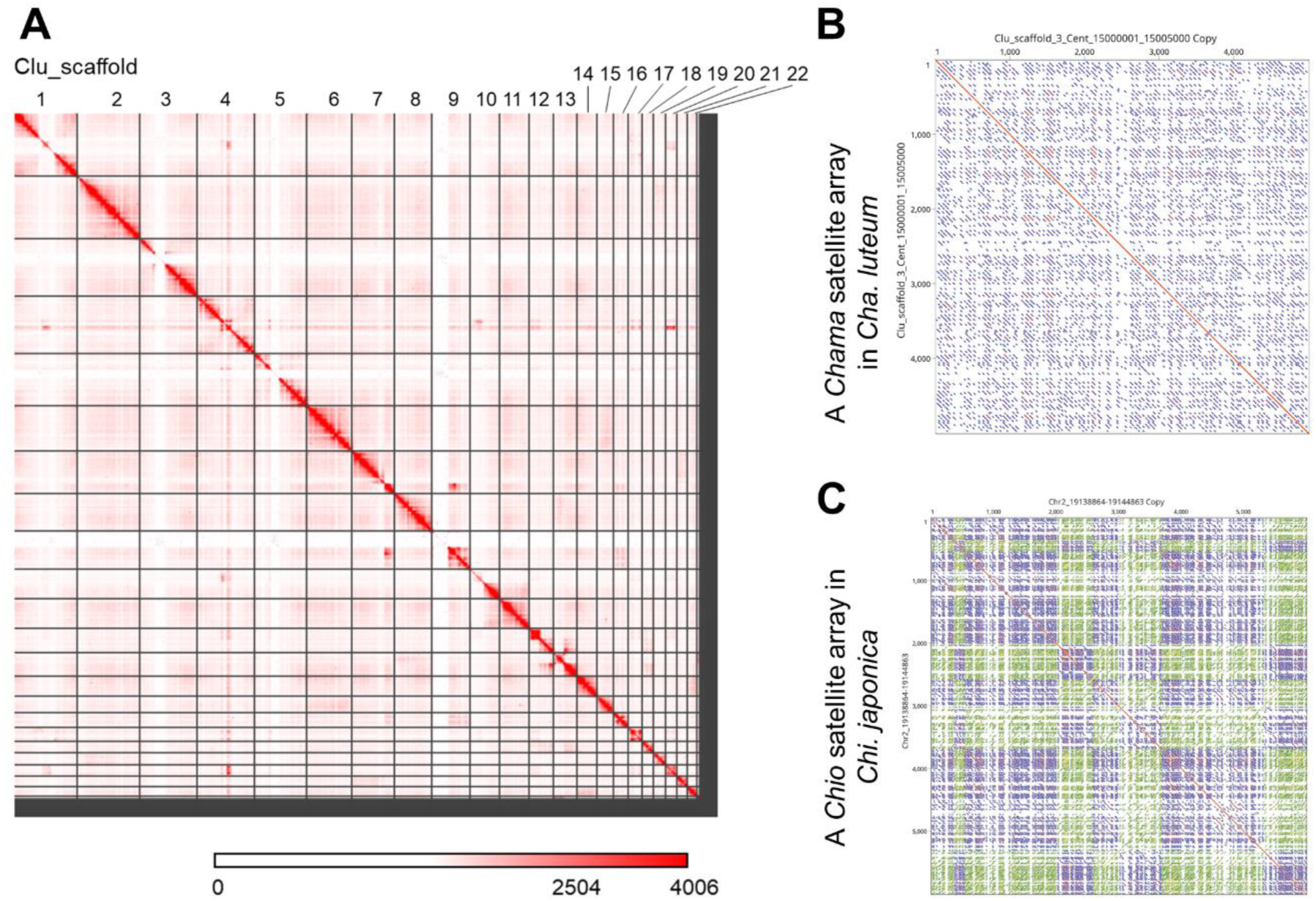
Hi-C map of *Cha. luteum* and contrasting centromere array organization in *Cha. luteum* and *Chi. japonica*. (A) Hi-C scaffolding of *Cha. luteum* contigs results in 22 scaffolds longer than 10 Mb (10–68 Mb) (Table S2). The color bar represents the contact frequencies, which are indicated by the number of links at a 5-Mb resolution. (B) The centromeric *Chama* monomers in *Cha. luteum* are arranged in the same orientation in a ∼5-kb satellite repeat array; while (C) in *Chi. japonica*, the orientation in a ∼6-kb centromeric *Chio* satellite array is different. Blue and green lines represent forward- and reverse-strand similarity, respectively. Red lines indicate regions of 100% sequence identity.

**Figure S3.**
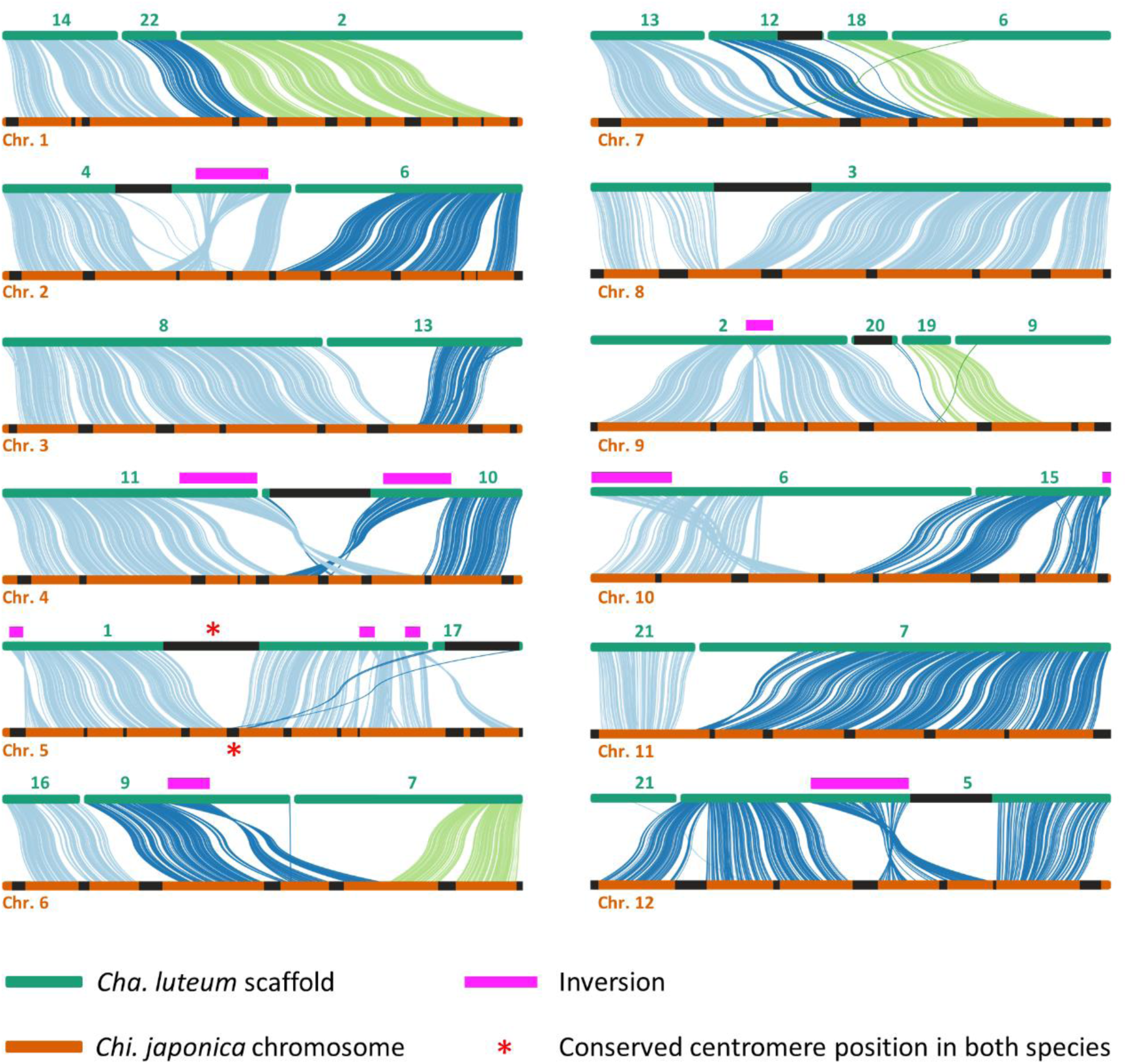
Holocentric *Chi. japonica* and monocentric *Cha. luteum* share broad-scale genome synteny, except for their centromeres. Visualization of syntenic orthologs of coding genes in the assembled top 22 scaffolds of *Cha. luteum* (green) and the 12 chromosome-level pseudomolecules of *Chi. japonica* (orange). The chromosome-level arrangement of orthologs between scaffold 3 of *Cha. luteum* and chromosome 8 of *Chi. japonica* is identical. In addition to chromosome-sized syntenic regions 11 large-scale inversions (magenta lines) and four inter-chromosomal translocations (scaffolds 2, 6, 7, and 13 of *Cha. luteum*) were identified. The conserved centromere positions in both species are marked by asterisks.

**Figure S4.**
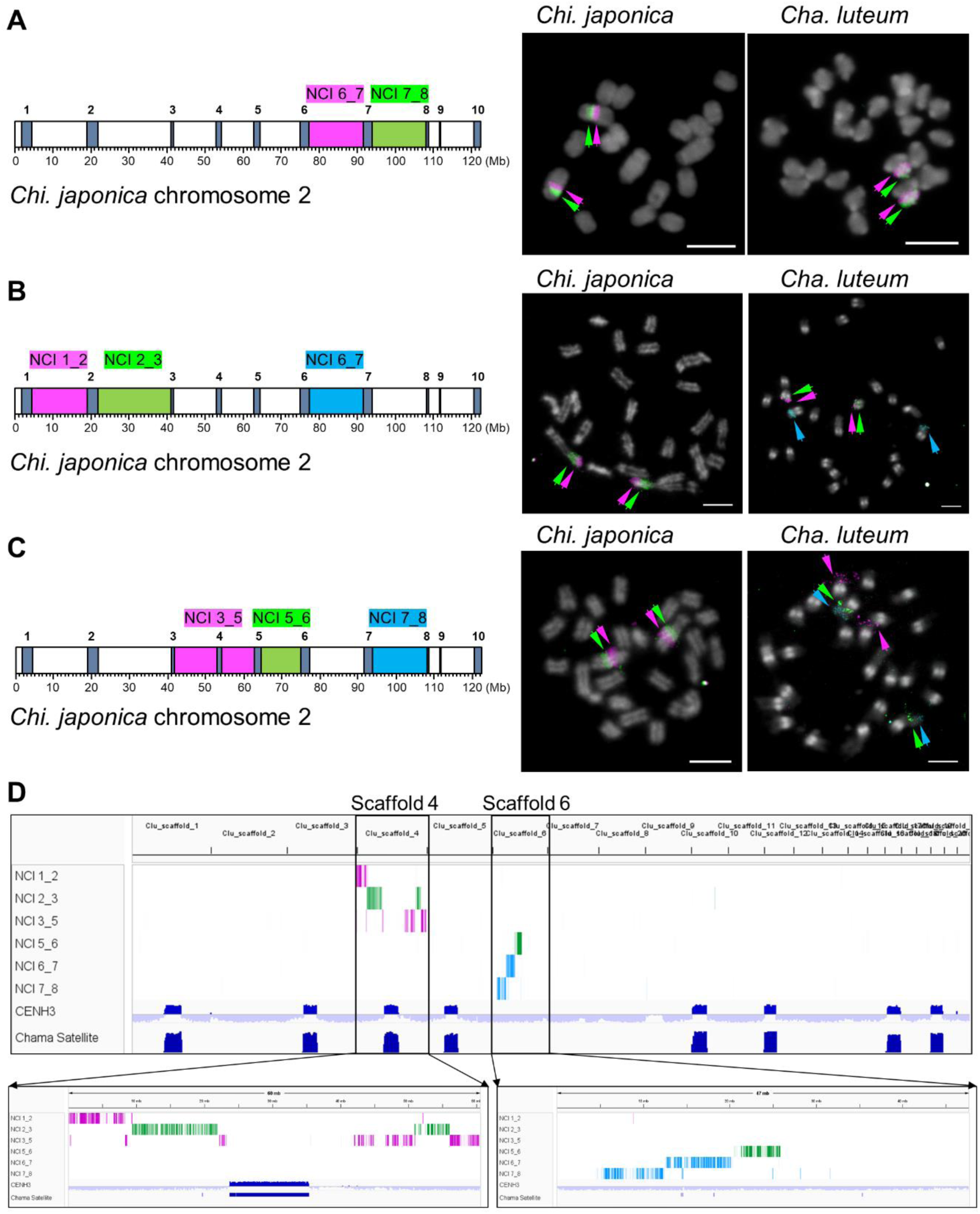
Comparative FISH mapping of the *Chi. japonica* chromosome 2-specific oligo-FISH probes on mitotic chromosomes of *Chi. japonica* and *Cha. luteum*. (A) Application of the multicolor non-centromeric interval (NCI)-specific oligo-FISH painting probes NCI 6_7 (magenta) and NCI 7_8 (green). (B) Application of the oligo painting probes NCI 1_2 (magenta), NCI 2_3 (green), and NCI 6_7 (blue). (C) Application of the oligo painting probes NCI 3_5 (magenta), NCI 5_6 (green), and NCI 7_8 (blue). Schemata show the chromosomal position and color of the probes along *Chi. japonica* chromosome 2. Colored arrows indicate the signals of the corresponding probes shown in the schemata. (D) *In silico* mapping of the six *Chi. japonica* chromosome 2-specific oligo probes onto the scaffolds 4 and 6 of *Cha. luteum*, which correspond to the three chromosome arms of two *Cha. luteum* chromosome pairs. Chromosomes were counterstained with DAPI. Scale bars = 5 µm.

**Figure S5.**
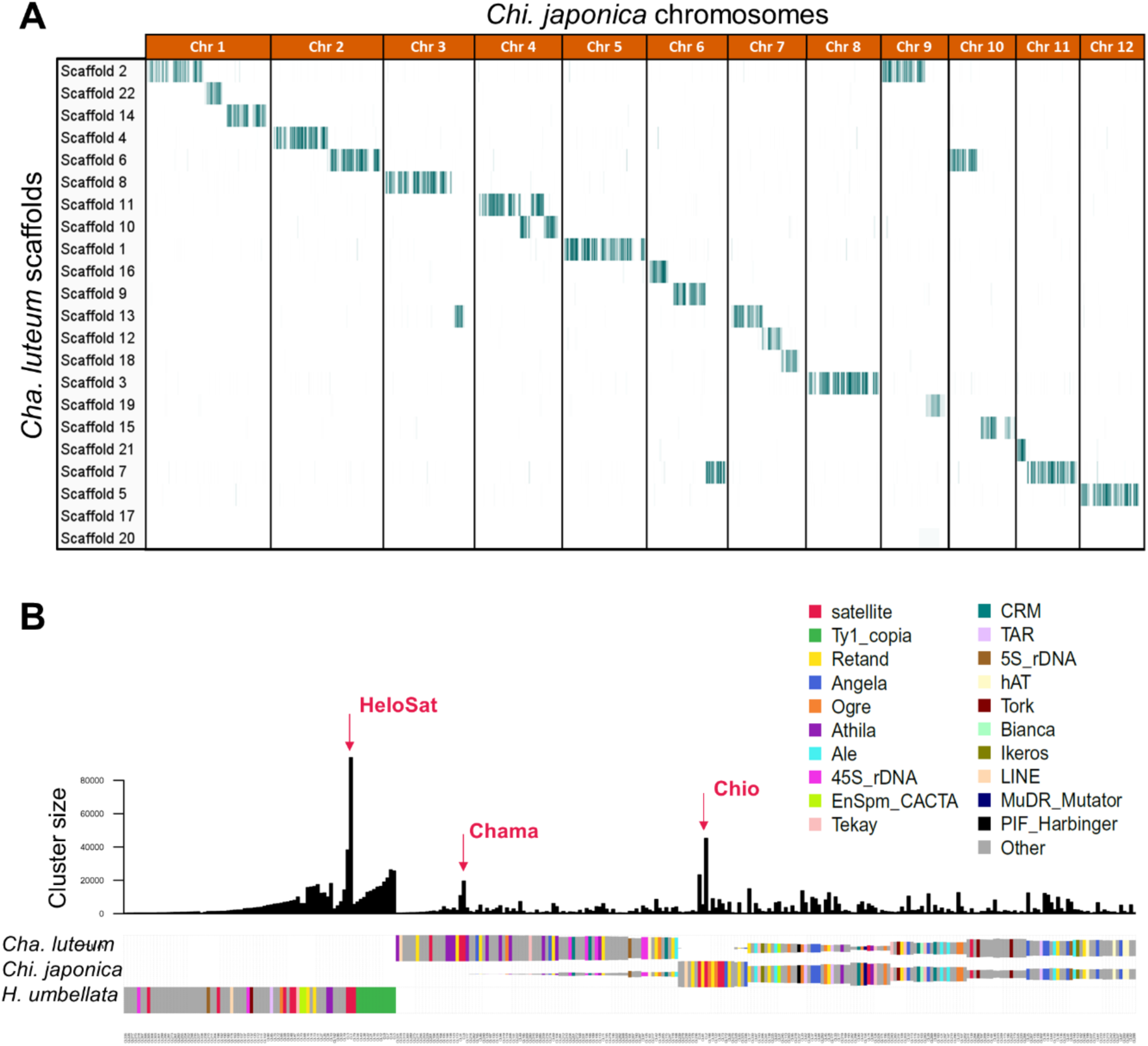
Comparative sequence analyses among Melanthiaceae species. (A) *In silico* comparative mapping of 12 *Chi. japonica*-based chromosome-wide oligo pools to the top 22 scaffolds of *Cha. luteum* confirmed high chromosomal collinearity between the two genomes and revealed no evidence of chromosome duplication. (B) Comparative repeat analysis using RepeatExplorer revealed no shared high-copy repeats between *Heloniopsis umbellata* and either *Cha. luteum* or *Chi. japonica.* Arrows indicate the position of the known, highly abundant satellite repeats *HeloSat* ^94^, *Chama* (this study) and *Chio* ^31^. The bar plot represents the abundance of each cluster. The annotation of clusters is shown in different colors.

**Figure S6.**
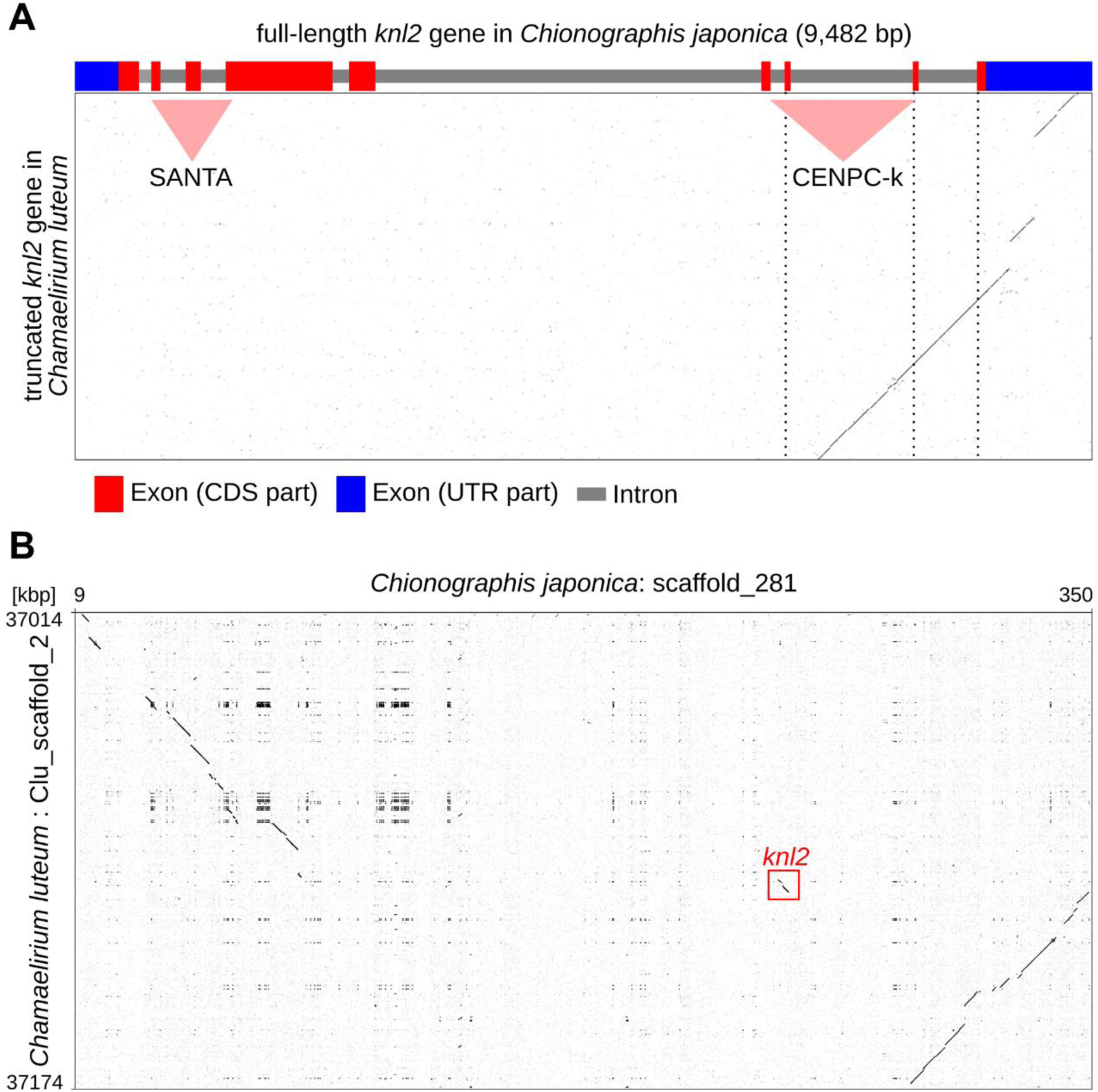
Loss of the *KNL2* gene in *Cha. luteum*. (A) Detailed dot plot comparison of the *KNL2* genes shows that most of the *KNL2* gene has been lost in *Cha. luteum*. Light red triangles mark the positions encoding the SANTA domain and the CENPC-k motif, indicating a complete loss of the SANTA-coding domain. Although a portion of the CENPC-k coding domain is retained in the truncated gene, it corresponds to the last nine amino acids at the C-terminus and could not be translated into a protein due to the absence of a start codon in the upstream region. (B) Dot plot comparison of orthologous loci between *Cha. luteum* and *Chi. japonica*. The position of the *KNL2* gene is marked by the red rectangle.

**Figure S7.**
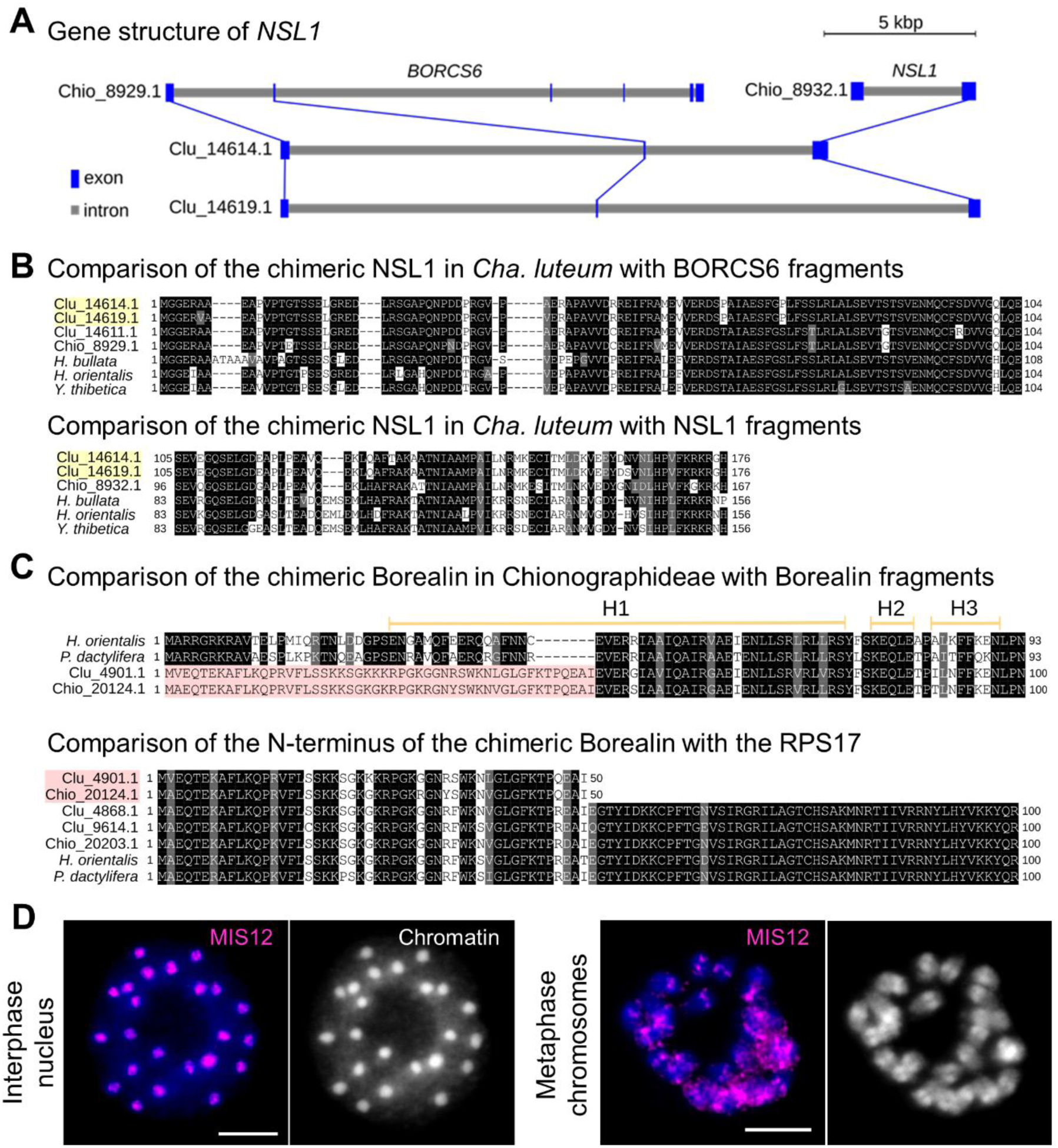
Chimeric origin of NSL1 in *Cha. luteum* and Borealin in both *Cha. luteum* and *Chi. japonica* and immunodetection of MIS12 in *Cha. luteum.* (A) In *Cha. luteum*, the *NSL1* gene has undergone significant changes through recombination with the *BORCS6* gene. The changes in these genes are shown by comparison with full-length homologous genes, *NSL1* (Chio_8932.1) and *BORCS6* (Chio_8929.1), identified in *Chi. japonica*. Note that *Cha. luteum* has two copies of the recombined *NSL1* gene (Clu_14614.1 and Clu_14619.1). (B) Comparison of the N- and C-termini of the chimeric NSL1 protein sequences in *Cha. luteum* with the corresponding parts of BORCS6 and NSL1 homologous proteins in *Chi. japonica* and three other Melanthiaceae species (*Helonias bullata, Heloniopsis orientalis*, and *Ypsilandra thibetica*). The recombinant NSL1 protein sequences in *Cha. luteum* are highlighted in yellow. (C) The N-terminus of Borealin in both *Cha. luteum* and *Chi. japonica* is replaced by the N-terminal fragment of ribosomal protein S17 (RPS17). Alignments of the protein sequences at the N-termini of the recombinant Borealin proteins with the corresponding parts of the intact Borealin and RPS17 in *H. orientalis* and *Phoenix dactylifera*. (D) Recruitment of MIS12 to centromeres of *Cha. luteum* remains unaffected. Immunolabelling of MIS12 in an interphase nucleus and on metaphase chromosomes of *Cha. luteum* using the *Chi. japonica*-specific anti-MIS12 antibody. Chromatin was counterstained with DAPI. Scale bars = 5 µm.

**Figure S8.**
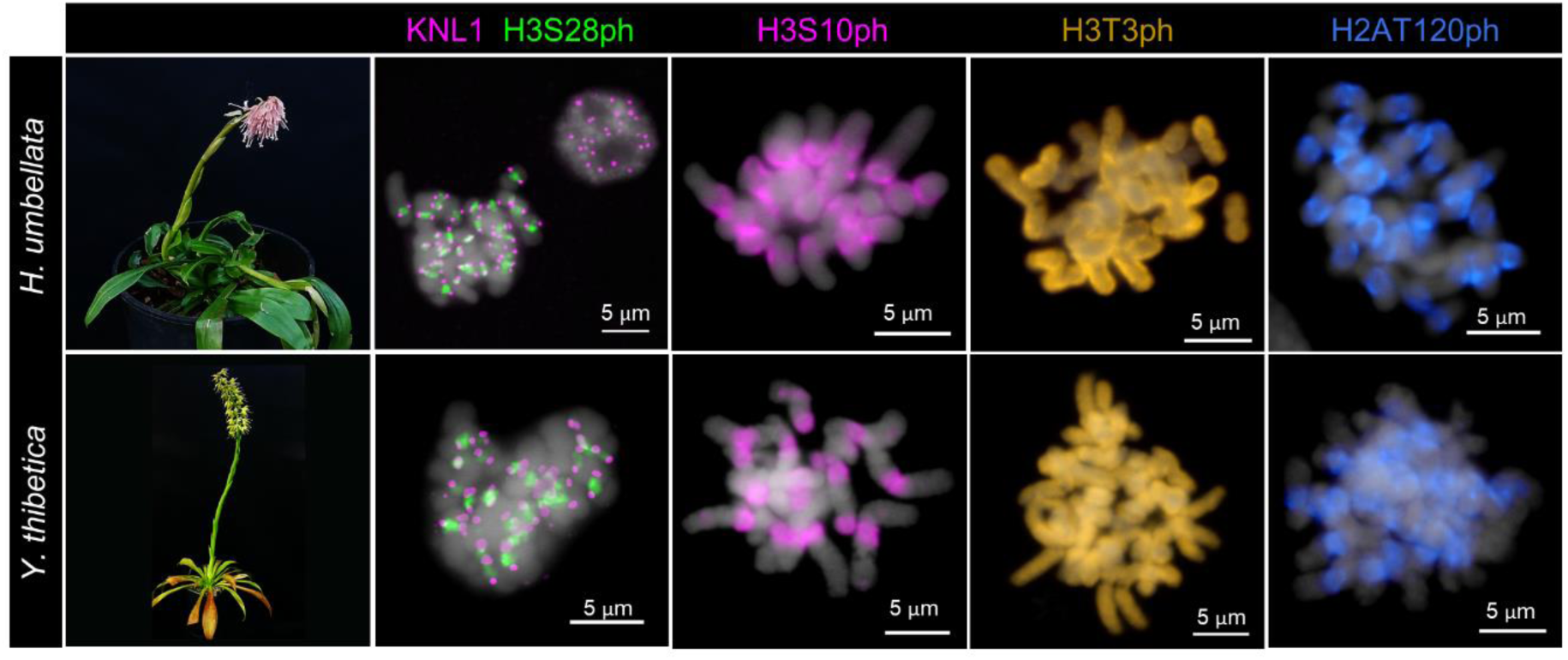
The Heloniadeae species possess monocentromeres with monocentric-typical histone phosphorylation patterns. Flowering *Heloniopsis umbellata* and *Ypsilandra thibetica* plants. Immunostaining of mitotic chromosomes using the kinetochore antibody KNL1 (magenta) and the cell cycle-dependent histone marks H3S28ph (green), H3S10ph (magenta), H3T3ph (yellow), and H2AT120ph (blue) in both species. Chromatin was counterstained with DAPI. Scale bars = 5 µm.

## Supplemental Information Document S2: Tables S1–S6

**Table S1.**
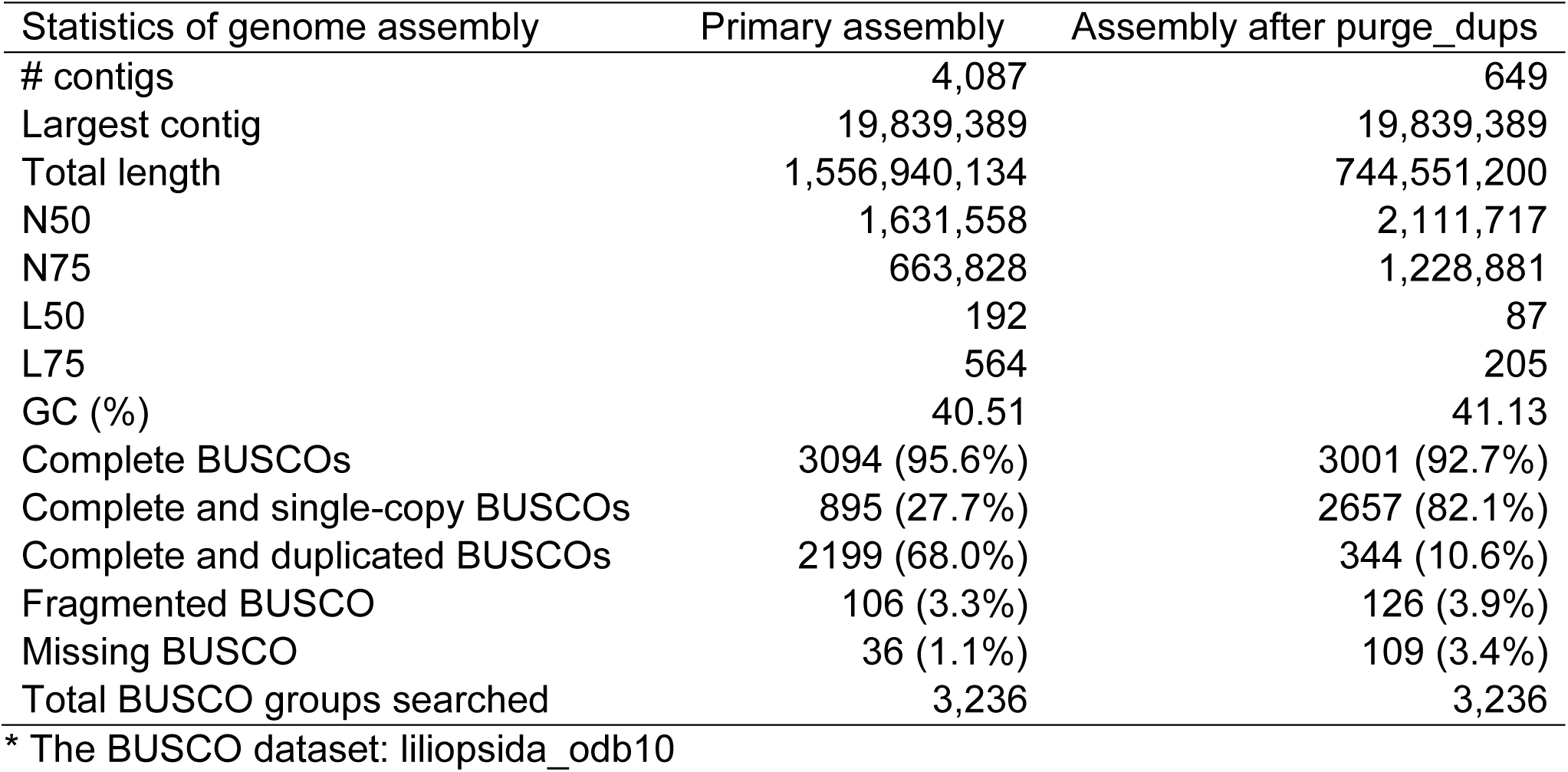
Genome assembly of *Cha. luteum*.

**Table S2.**
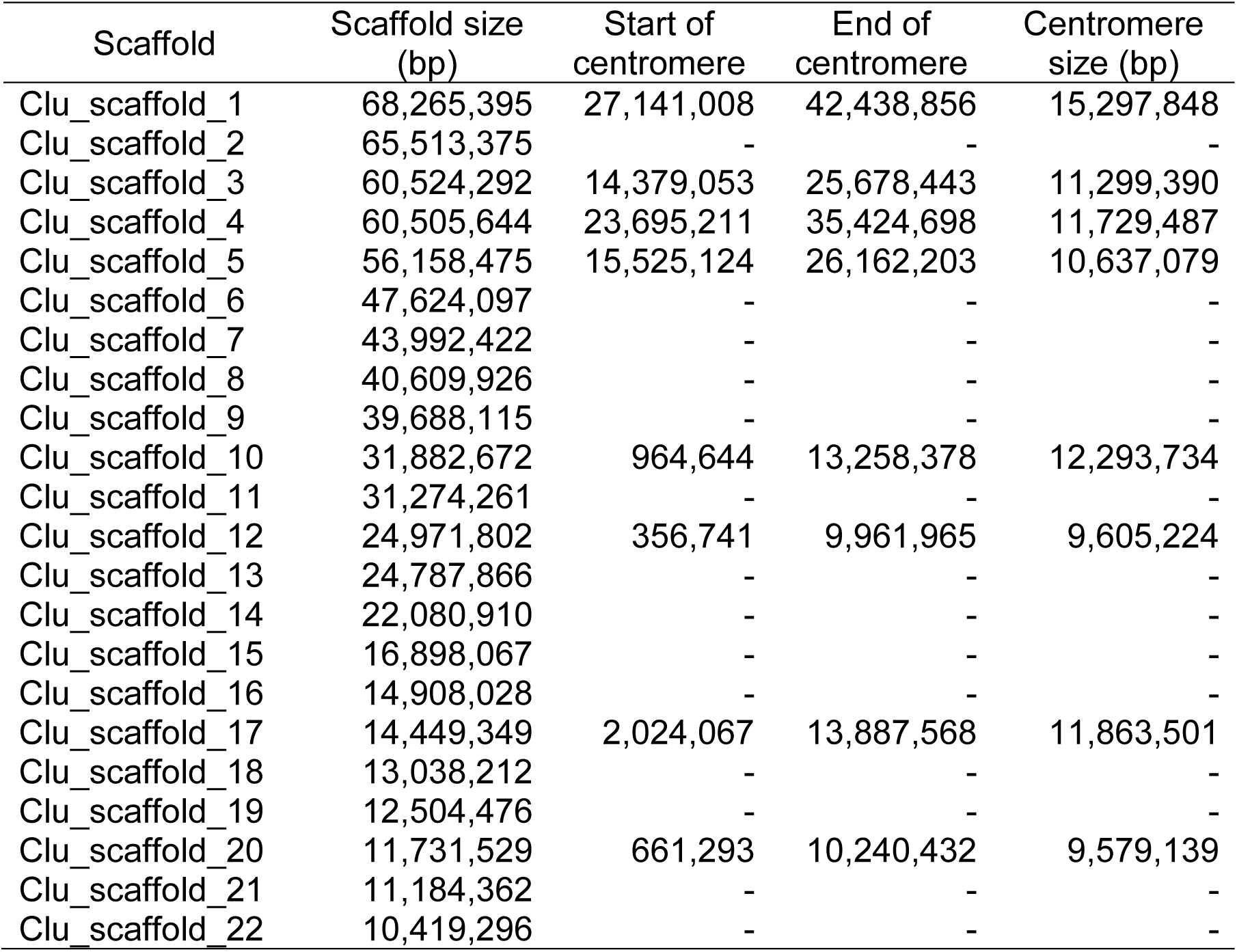
Hi-C scaffolding and CENH3-ChIPseq determined centromere position and size on the top 22 scaffolds of *Cha. luteum*.

| Scaffold | Scaffold size (bp) | Start of centromere | End of centromere | Centromere size (bp) |
| --- | --- | --- | --- | --- |
| Clu_scaffold_1 | 68,265,395 | 27,141,008 | 42,438,856 | 15,297,848 |
| Clu_scaffold_2 | 65,513,375 | - | - | - |
| Clu_scaffold_3 | 60,524,292 | 14,379,053 | 25,678,443 | 11,299,390 |
| Clu_scaffold_4 | 60,505,644 | 23,695,211 | 35,424,698 | 11,729,487 |
| Clu_scaffold_5 | 56,158,475 | 15,525,124 | 26,162,203 | 10,637,079 |
| Clu_scaffold_6 | 47,624,097 | - | - | - |
| Clu_scaffold_7 | 43,992,422 | - | - | - |
| Clu_scaffold_8 | 40,609,926 | - | - | - |
| Clu_scaffold_9 | 39,688,115 | - | - | - |
| Clu_scaffold_10 | 31,882,672 | 964,644 | 13,258,378 | 12,293,734 |
| Clu_scaffold_11 | 31,274,261 | - | - | - |
| Clu_scaffold_12 | 24,971,802 | 356,741 | 9,961,965 | 9,605,224 |
| Clu_scaffold_13 | 24,787,866 | - | - | - |
| Clu_scaffold_14 | 22,080,910 | - | - | - |
| Clu_scaffold_15 | 16,898,067 | - | - | - |
| Clu_scaffold_16 | 14,908,028 | - | - | - |
| Clu_scaffold_17 | 14,449,349 | 2,024,067 | 13,887,568 | 11,863,501 |
| Clu_scaffold_18 | 13,038,212 | - | - | - |
| Clu_scaffold_19 | 12,504,476 | - | - | - |
| Clu_scaffold_20 | 11,731,529 | 661,293 | 10,240,432 | 9,579,139 |
| Clu_scaffold_21 | 11,184,362 | - | - | - |
| Clu_scaffold_22 | 10,419,296 | - | - | - |

**Table S3.**
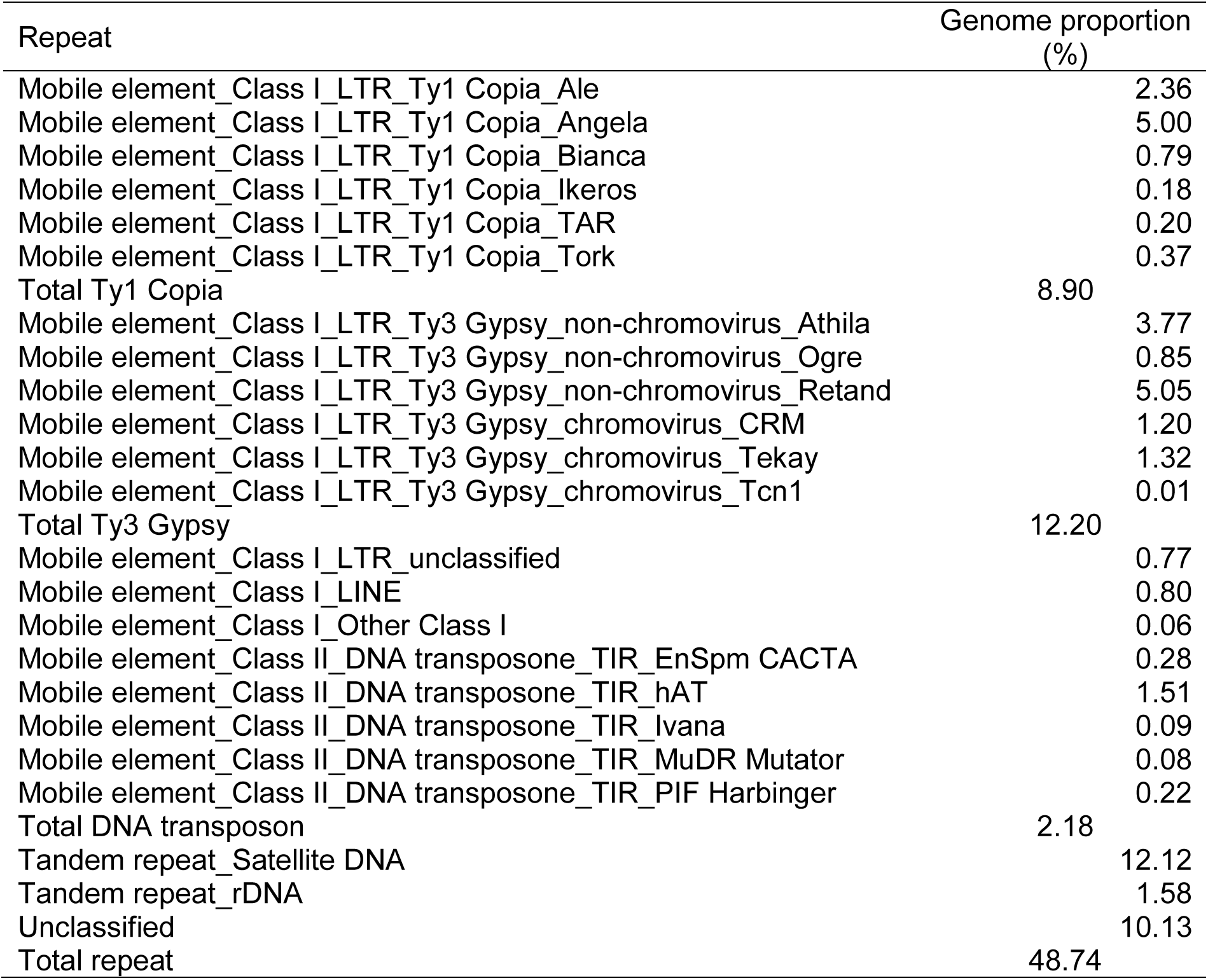
Repetitive composition of the *Cha. luteum* genome analyzed using RepeatExplorer.

**Table S4.** Centromere proportion in different monocentric species.

| Species | Genome size (Mb/1C) | Chromosome number (n) | Average chromosome size (Mb) | Average centromere size (Mb)* | Average size ratio of centromere/ chromosome | Reference |
| --- | --- | --- | --- | --- | --- | --- |
| *Chamaelirium luteum* | 887.4 | 12 | 74.0 | 11.5 | 15.6% | In this study |
| *Arabidopsis thaliana* | 156.5 | 5 | 31.3 | 2.5 | 8.1% | (Naish et al., 2021)<sup>1</sup> |
| *Brachypodium hybridum* | 616.1 | 15 | 41.1 | 0.9 | 2.1% | (Chen et al., 2024)<sup>2</sup> |
| *Brachypodium distachyon* | 313.0 | 5 | 62.6 | 0.6 | 1.0% | (Chen et al., 2024)<sup>2</sup> |
| *Brachypodium stacei* | 273.8 | 10 | 27.4 | 1.1 | 4.0% | (Chen et al., 2024)<sup>2</sup> |
| *Brassica rapa* (Chiifu v4.0) | 782.4 | 10 | 78.2 | 6.9 | 8.8% | (Zhang et al., 2023)<sup>3</sup> |
| *Capsicum annuum* | 3090.5 | 12 | 257.5 | 2.5 | 1.0% | (Chen et al., 2024)<sup>4</sup> |
| *Chlorella pyrenoidosa* | 53.4 | 13 | 4.1 | 0.0 | 1.0% | (Wang et al., 2024)<sup>5</sup> |
| *Chlorella sorokiniana* | 58.1 | 12 | 4.8 | 0.1 | 1.2% | (Wang et al., 2024)<sup>5</sup> |
| *Citrus reticulata* | 459.7 | 9 | 51.1 | 2.1 | 4.1% | (Xia et al., 2020)<sup>6</sup> |
| *Drosophila melanogaster* | 175.0 | 4 | 43.8 | 0.5 | 1.1% | (Chang et al., 2019)<sup>7</sup> |
| *Forsythia suspensa* | 688.8 | 14 | 49.2 | 0.9 | 1.9% | (Cui et al., 2024)<sup>8</sup> |
| *Glycine max* ZH13 | 1105.1 | 20 | 55.3 | 1.3 | 2.4% | (Liu et al., 2023)<sup>9</sup> |
| *Gossypium raimondii* | 880.2 | 13 | 67.7 | 1.9 | 2.7% | (Huang et al., 2024)<sup>10</sup> |
| *Gossypium anomalum* | 1359.4 | 13 | 104.6 | 1.1 | 1.1% | (Ding et al., 2023)<sup>11</sup> |
| *Homo sapiens* | 3178.5 | 23 | 138.2 | 5.1 | 3.7% | (Logsdon et al., 2024)<sup>12</sup> |
| *Juncus effusus* | 270.4 | 21 | 12.9 | 0.4 | 3.3% | (Dias et al., 2024)<sup>13</sup> |
| *Lactuca sativa* | 3237.2 | 9 | 359.7 | 3.4 | 1.0% | (Wang et al., 2024)<sup>14</sup> |
| *Nicotiana benthamiana* | 3129.6 | 19 | 164.7 | 2.0 | 1.2% | (Chen et al., 2024)<sup>15</sup> |
| *Oryza sativa* | 489.0 | 12 | 40.8 | 1.4 | 3.4% | (Song et al., 2021)<sup>16</sup> |
| *Secale cereale* | 8606.4 | 7 | 1229.5 | 8.6 | 0.7% | (Liu et al., 2024)<sup>17</sup> |
| *Solanum tuberosum* | 870.4 | 12 | 72.5 | 0.9 | 1.2% | (Pham et al., 2020)<sup>18</sup> |
| *Triticum monococcum* | 6063.6 | 14 | 433.1 | 5.5 | 1.3% | (Ahmed et al., 2023)<sup>19</sup> |
| *Triticum aestivum* | 16919.4 | 21 | 805.7 | 7.9 | 1.0% | (Su et al., 2019)<sup>20</sup> |
| *Vitis vinifera* | 391.2 | 19 | 20.6 | 1.4 | 6.8% | (Shi et al., 2023)<sup>21</sup> |
| *Zea mays* Mo17 | 2640.6 | 10 | 264.1 | 2.2 | 0.8% | (Chen et al., 2023)<sup>22</sup> |

\*Centromere size was determined based on the CENH3-ChIP-seq results in the corresponding reference

**Table S5.**
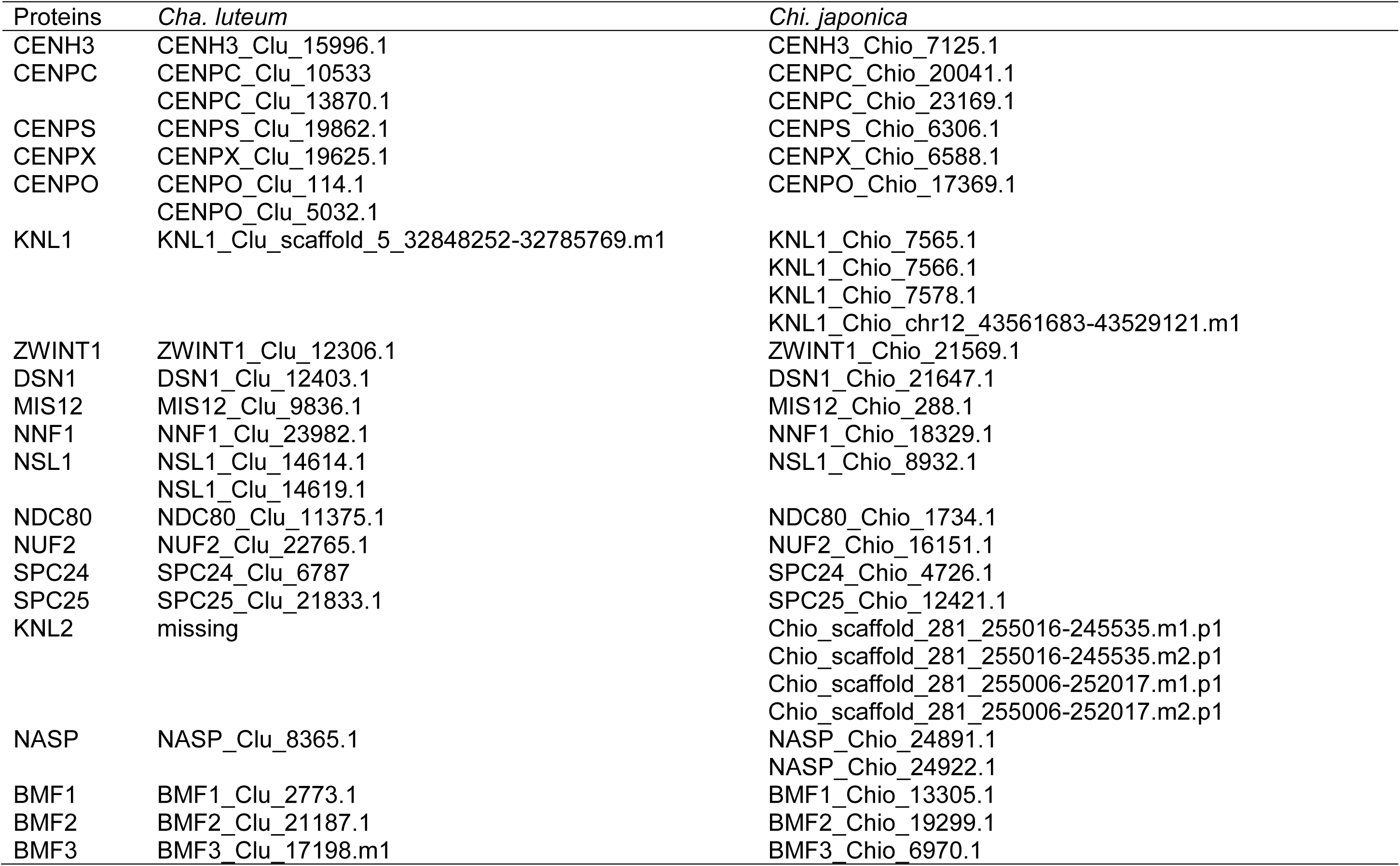
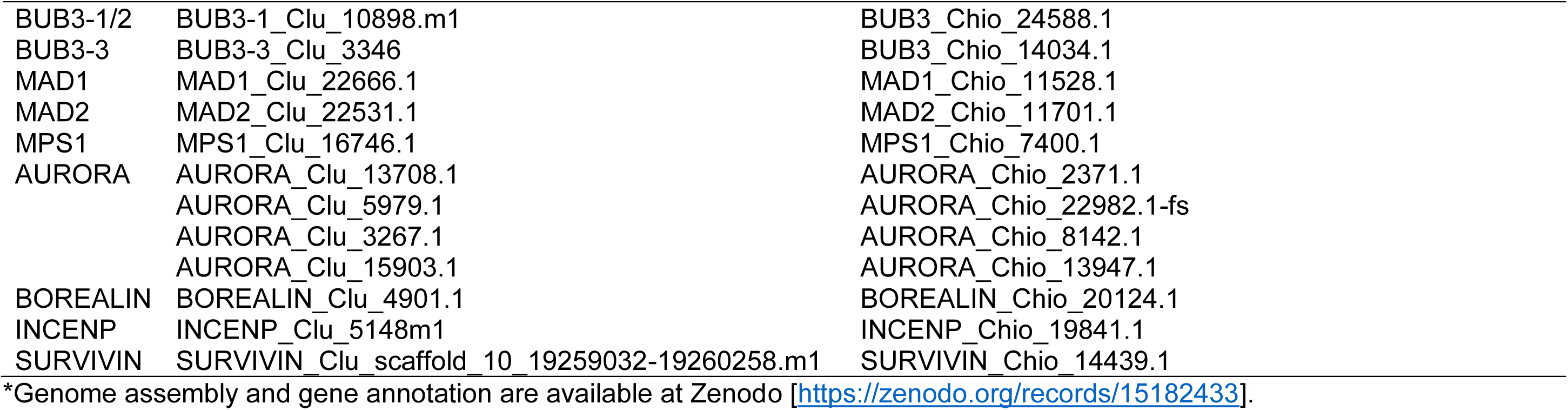
Sequence IDs of the analyzed kinetochore proteins in the gene annotation of *Cha. luteum* and *Chi. japonica*.

| Proteins | *Cha. luteum* | *Chi. japonica* |
| --- | --- | --- |
| CENH3 | CENH3_Clu_15996.1 | CENH3_Chio_7125.1 |
| CENPC | CENPC_Clu_10533 | CENPC_Chio_20041.1 |
|  | CENPC_Clu_13870.1 | CENPC_Chio_23169.1 |
| CENPS | CENPS_Clu_19862.1 | CENPS_Chio_6306.1 |
| CENPX | CENPX_Clu_19625.1 | CENPX_Chio_6588.1 |
| CENPO | CENPO_Clu_114.1 | CENPO_Chio_17369.1 |
|  | CENPO_Clu_5032.1 |  |
| KNL1 | KNL1_Clu_scaffold_5_32848252-32785769.m1 | KNL1_Chio_7565.1 |
|  |  | KNL1_Chio_7566.1 |
|  |  | KNL1_Chio_7578.1 |
|  |  | KNL1_Chio_chr12_43561683-43529121.m1 |
| ZWINT1 | ZWINT1_Clu_12306.1 | ZWINT1_Chio_21569.1 |
| DSN1 | DSN1_Clu_12403.1 | DSN1_Chio_21647.1 |
| MIS12 | MIS12_Clu_9836.1 | MIS12_Chio_288.1 |
| NNF1 | NNF1_Clu_23982.1 | NNF1_Chio_18329.1 |
| NSL1 | NSL1_Clu_14614.1 | NSL1_Chio_8932.1 |
|  | NSL1_Clu_14619.1 |  |
| NDC80 | NDC80_Clu_11375.1 | NDC80_Chio_1734.1 |
| NUF2 | NUF2_Clu_22765.1 | NUF2_Chio_16151.1 |
| SPC24 | SPC24_Clu_6787 | SPC24_Chio_4726.1 |
| SPC25 | SPC25_Clu_21833.1 | SPC25_Chio_12421.1 |
| KNL2 | missing | Chio_scaffold_281_255016-245535.m1.p1 |
|  |  | Chio_scaffold_281_255016-245535.m2.p1 |
|  |  | Chio_scaffold_281_255006-252017.m1.p1 |
|  |  | Chio_scaffold_281_255006-252017.m2.p1 |
| NASP | NASP_Clu_8365.1 | NASP_Chio_24891.1 |
|  |  | NASP_Chio_24922.1 |
| BMF1 | BMF1_Clu_2773.1 | BMF1_Chio_13305.1 |
| BMF2 | BMF2_Clu_21187.1 | BMF2_Chio_19299.1 |
| BMF3 | BMF3_Clu_17198.m1 | BMF3_Chio_6970.1 |
| BUB3-1/2 | BUB3-1_Clu_10898.m1 | BUB3_Chio_24588.1 |
| BUB3-3 | BUB3-3_Clu_3346 | BUB3_Chio_14034.1 |
| MAD1 | MAD1_Clu_22666.1 | MAD1_Chio_11528.1 |
| MAD2 | MAD2_Clu_22531.1 | MAD2_Chio_11701.1 |
| MPS1 | MPS1_Clu_16746.1 | MPS1_Chio_7400.1 |
| AURORA | AURORA_Clu_13708.1 | AURORA_Chio_2371.1 |
|  | AURORA_Clu_5979.1 | AURORA_Chio_22982.1-fs |
|  | AURORA_Clu_3267.1 | AURORA_Chio_8142.1 |
|  | AURORA_Clu_15903.1 | AURORA_Chio_13947.1 |
| BOREALIN | BOREALIN_Clu_4901.1 | BOREALIN_Chio_20124.1 |
| INCENP | INCENP_Clu_5148m1 | INCENP_Chio_19841.1 |
| SURVIVIN | SURVIVIN_Clu_scaffold_10_19259032-19260258.m1 | SURVIVIN_Chio_14439.1 |

\*Genome assembly and gene annotation are available at Zenodo [https://zenodo.org/records/15182433].

**Table S6.**
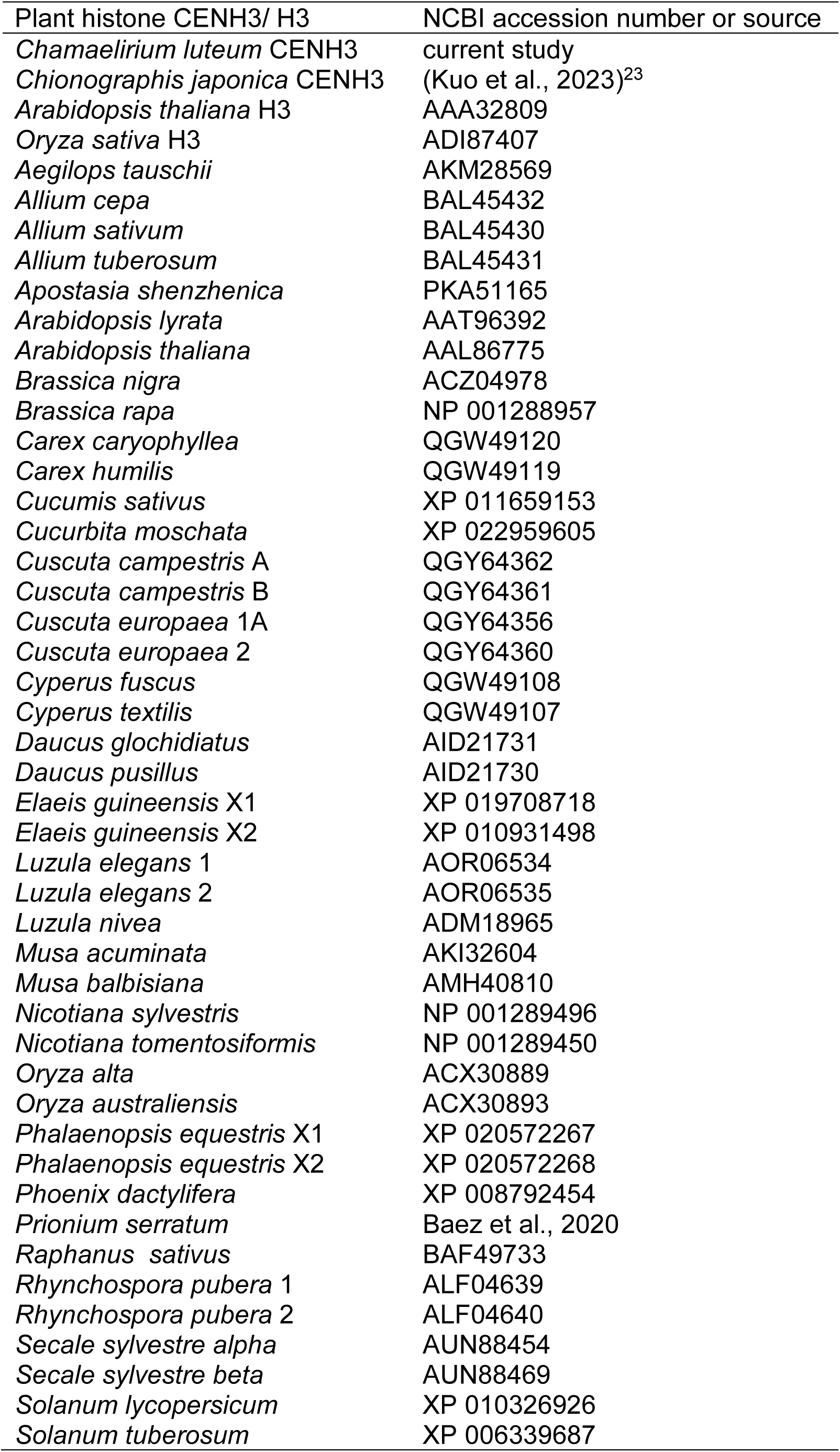
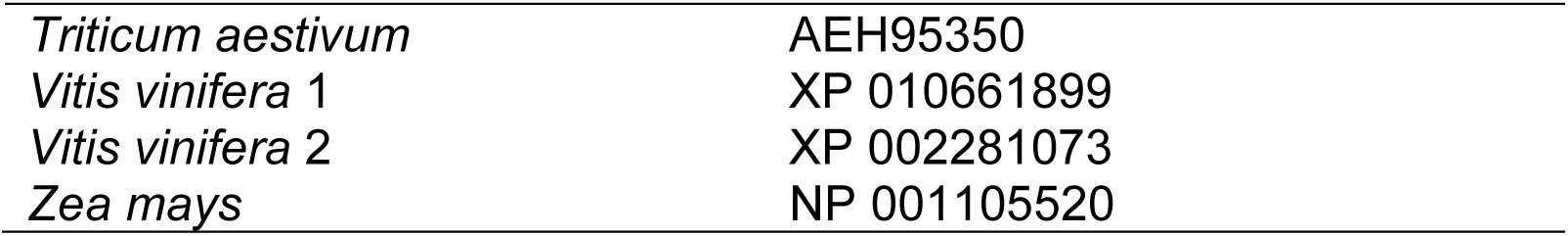
NCBI accession numbers and sources of CENH3 and H3 protein sequences.

## Notes

### Competing Interest Statement

The authors have declared no competing interest.

## References

1. Flemming, W. (1882). Zellsubstanz, Kern und Zelltheilung. Book Verlag F. C. W. Vogel, Leipzig. https://books.google.de/books?hl=de&lr=&id=J4RIAAAAYAAJ&oi=fnd&pg=PA1&ots=MxJ8m5uJWo&sig=l6y_oK8gWaHaxroQRKlsMlk30YI&redir_esc=y#v=onepage&q&f=false.

2. Schubert, V., Neumann, P., Marques, A., Heckmann, S., Macas, J., Pedrosa-Harand, A., Schubert, I., Jang, T.S., and Houben, A. (2020). Super-Resolution Microscopy Reveals Diversity of Plant Centromere Architecture. Int J Mol Sci 21. 10.3390/ijms21103488.

3. Kuo, Y.T., Schubert, V., Marques, A., Schubert, I., and Houben, A. (2024). Centromere diversity: How different repeat-based holocentromeres may have evolved. Bioessays, e2400013. 10.1002/bies.202400013.

4. Macas, J., Avila Robledillo, L., Kreplak, J., Novak, P., Koblizkova, A., Vrbova, I., Burstin, J., and Neumann, P. (2023). Assembly of the 81.6 Mb centromere of pea chromosome 6 elucidates the structure and evolution of metapolycentric chromosomes. PLoS Genet 19, e1010633. 10.1371/journal.pgen.1010633.

5. Neumann, P., Navratilova, A., Schroeder-Reiter, E., Koblizkova, A., Steinbauerova, V., Chocholova, E., Novak, P., Wanner, G., and Macas, J. (2012). Stretching the rules: monocentric chromosomes with multiple centromere domains. Plos Genet 8, e1002777. 10.1371/journal.pgen.1002777.

6. Grzan, T., Despot-Slade, E., Mestrovic, N., Plohl, M., and Mravinac, B. (2020). CenH3 distribution reveals extended centromeres in the model beetle Tribolium castaneum. Plos Genet 16, e1009115. 10.1371/journal.pgen.1009115.

7. McKinley, K.L., and Cheeseman, I.M. (2016). The molecular basis for centromere identity and function. Nat Rev Mol Cell Biol 17, 16–29. 10.1038/nrm.2015.5.

8. Ochs, F., Green, C., Szczurek, A.T., Pytowski, L., Kolesnikova, S., Brown, J., Gerlich, D.W., Buckle, V., Schermelleh, L., and Nasmyth, K.A. (2024). Sister chromatid cohesion is mediated by individual cohesin complexes. Science 383, 1122–1130. 10.1126/science.adl4606.

9. Schmitz, M.L., Higgins, J.M.G., and Seibert, M. (2020). Priming chromatin for segregation: functional roles of mitotic histone modifications. Cell Cycle 19, 625–641. 10.1080/15384101.2020.1719585.

10. Houben, A., Wako, T., Furushima-Shimogawara, R., Presting, G., Kunzel, G., Schubert, I., and Fukui, K. (1999). The cell cycle dependent phosphorylation of histone H3 is correlated with the condensation of plant mitotic chromosomes. Plant Journal 18, 675–679.

11. Gernand, D., Demidov, D., and Houben, A. (2003). The temporal and spatial pattern of histone H3 phosphorylation at serine 28 and serine 10 is similar in plants but differs between mono- and polycentric chromosomes. Cytogenet Genome Res 101, 172–176.

12. Caperta, A.D., Rosa, M., Delgado, M., Karimi, R., Demidov, D., Viegas, W., and Houben, A. (2008). Distribution patterns of phosphorylated Thr 3 and Thr 32 of histone H3 in plant mitosis and meiosis. Cytogenet Genome Res 122, 73–79. 10.1159/000151319.

13. Agueci, F., Karimi, R., and Houben, A. (2007). Characterisation of nima- and haspin-like kinases in arabidopsis thaliana. Chromosome Research 15, 98–98.

14. Dong, Q., and Han, F. (2012). Phosphorylation of histone H2A is associated with centromere function and maintenance in meiosis. Plant J 71, 800–809. 10.1111/j.1365-313X.2012.05029.x.

15. Demidov, D., Schubert, V., Kumke, K., Weiss, O., Karimi-Ashtiyani, R., Buttlar, J., Heckmann, S., Wanner, G., Dong, Q., Han, F., and Houben, A. (2014). Anti-phosphorylated histone H2AThr120: a universal microscopic marker for centromeric chromatin of mono- and holocentric plant species. Cytogenet Genome Res 143, 150–156. 10.1159/000360018.

16. Marques, A., and Drinnenberg, I.A. (2025). Same but different: Centromere regulations in holocentric insects and plants. Curr Opin Cell Biol 93, 102484. 10.1016/j.ceb.2025.102484.

17. Senaratne, A.P., Cortes-Silva, N., and Drinnenberg, I.A. (2022). Evolution of holocentric chromosomes: Drivers, diversity, and deterrents. Semin Cell Dev Biol. 10.1016/j.semcdb.2022.01.003.

18. Drinnenberg, I.A., Henikoff, S., and Malik, H.S. (2016). Evolutionary Turnover of Kinetochore Proteins: A Ship of Theseus? Trends Cell Biol 26, 498–510. 10.1016/j.tcb.2016.01.005.

19. Neumann, P., Oliveira, L., Jang, T.S., Novak, P., Koblizkova, A., Schubert, V., Houben, A., and Macas, J. (2023). Disruption of the standard kinetochore in holocentric Cuscuta species. Proc Natl Acad Sci U S A 120, e2300877120. 10.1073/pnas.2300877120.

20. Drinnenberg, I.A., deYoung, D., Henikoff, S., and Malik, H.S. (2014). Recurrent loss of CenH3 is associated with independent transitions to holocentricity in insects. Elife 3. 10.7554/eLife.03676.

21. Senaratne, A.P., Muller, H., Fryer, K.A., Kawamoto, M., Katsuma, S., and Drinnenberg, I.A. (2021). Formation of the CenH3-Deficient Holocentromere in Lepidoptera Avoids Active Chromatin. Curr Biol 31, 173–181 e177. 10.1016/j.cub.2020.09.078.

22. Cortes-Silva, N., Ulmer, J., Kiuchi, T., Hsieh, E., Cornilleau, G., Ladid, I., Dingli, F., Loew, D., Katsuma, S., and Drinnenberg, I.A. (2020). CenH3-Independent Kinetochore Assembly in Lepidoptera Requires CCAN, Including CENP-T. Current Biology 30, 561–572.e510. 10.1016/j.cub.2019.12.014.

23. Tanaka, N., and Tanaka, N. (1977). Chromosome studies in Chionographis (Liliaceae).1. Holokinetic nature of chromosomes in Chionographis japonica Maxim. Cytologia 42, 753–763.

24. Tanaka, N., and Tanaka, N. (1979). Chromosome-Studies in Chionographis (Liliaceae).2. Morphological-Characteristics of the Somatic Chromosomes of 4 Japanese Members. Cytologia 44, 935–949.

25. Tanaka, N. (2020). Chromosomal Variation in Populations of Chamaelirium hisauchianum (Melanthiaceae) with Holocentric Chromosomes. Cytologia 85, 115–122. 10.1508/cytologia.85.115.

26. Tanaka, N. (2020). High Stability in Chromosomal Traits of Chamaelirium japonicum and C. koidzuminum (Melanthiaceae) with Holocentric Chromosomes. Cytologia 85, 33–40. 10.1508/cytologia.85.33.

27. Pellicer, J., Kelly, L.J., Leitch, I.J., Zomlefer, W.B., and Fay, M.F. (2014). A universe of dwarfs and giants: genome size and chromosome evolution in the monocot family Melanthiaceae. New Phytol 201, 1484–1497. 10.1111/nph.12617.

28. Tanaka, N. (2020). Chromosomal traits of Chamaelirium luteum (Melanthiaceae) with particular focus on the large heterochromatic centromeres. Taiwania 65, 286–294. 10.6165/tai.2020.65.286.

29. Kim, C., Kim, S.-C., and Kim, J.-H. (2019). Historical Biogeography of Melanthiaceae: A Case of Out-of-North America Through the Bering Land Bridge. Frontiers in Plant Science 10. 10.3389/fpls.2019.00396.

30. Tanaka, N. (2017). A synopsis of the genus Chamaelirium (Melanthiaceae) with a new infrageneric classification including Chionographis.

31. Kuo, Y.-T., Câmara, A.S., Schubert, V., Neumann, P., Macas, J., Melzer, M., Chen, J., Fuchs, J., Abel, S., Klocke, E., et al. (2023). Holocentromeres can consist of merely a few megabase-sized satellite arrays. Nat Commun 14, 3502. 10.1038/s41467-023-38922-7.

32. Hofstatter, P.G., Thangavel, G., Lux, T., Neumann, P., Vondrak, T., Novak, P., Zhang, M., Costa, L., Castellani, M., and Scott, A. (2022). Repeat-based holocentromeres influence genome architecture and karyotype evolution. Cell 185, 3153–3168. e3118.

33. Talbert, P.B., and Henikoff, S. (2020). What makes a centromere? Exp Cell Res 389. 10.1016/j.yexcr.2020.111895.

34. Fransz, P., de Jong, J.H., Lysak, M., Castiglione, M.R., and Schubert, I. (2002). Interphase chromosomes in Arabidopsis are organized as well defined chromocenters from which euchromatin loops emanate. P Natl Acad Sci USA 99, 14584–14589. 10.1073/pnas.212325299.

35. Kuo, Y.-T., Ishii, T., Fuchs, J., Hsieh, W.-H., Houben, A., and Lin, Y.-R. (2021). The Evolutionary Dynamics of Repetitive DNA and Its Impact on the Genome Diversification in the Genus Sorghum. Frontiers in Plant Science 12. 10.3389/fpls.2021.729734.

36. Muller, H., Gil, J., and Drinnenberg, I.A. (2019). The Impact of Centromeres on Spatial Genome Architecture. Trends Genet 35, 565–578. 10.1016/j.tig.2019.05.003.

37. van Hooff, J.J., Tromer, E., van Wijk, L.M., Snel, B., and Kops, G.J. (2017). Evolutionary dynamics of the kinetochore network in eukaryotes as revealed by comparative genomics. EMBO Rep. 10.15252/embr.201744102.

38. Ishii, M., and Akiyoshi, B. (2022). Plasticity in centromere organization and kinetochore composition: Lessons from diversity. Curr Opin Cell Biol 74, 47–54. 10.1016/j.ceb.2021.12.007.

39. Petrovic, A., Pasqualato, S., Dube, P., Krenn, V., Santaguida, S., Cittaro, D., Monzani, S., Massimiliano, L., Keller, J., Tarricone, A., et al. (2010). The MIS12 complex is a protein interaction hub for outer kinetochore assembly. J Cell Biol 190, 835–852. 10.1083/jcb.201002070.

40. Petrovic, A., Keller, J., Liu, Y., Overlack, K., John, J., Dimitrova, Y.N., Jenni, S., van Gerwen, S., Stege, P., Wohlgemuth, S., et al. (2016). Structure of the MIS12 Complex and Molecular Basis of Its Interaction with CENP-C at Human Kinetochores. Cell 167, 1028–1040 e1015. 10.1016/j.cell.2016.10.005.

41. Oliveira, L., Neumann, P., Mata-Sucre, Y., Kuo, Y.-T., Marques, A., Schubert, V., and Macas, J. (2024). KNL1 and NDC80 represent new universal markers for the detection of functional centromeres in plants. Chromosome Research 32, 3. 10.1007/s10577-024-09747-x.

42. Houben, A., Demidov, D., Caperta, A.D., Karimi, R., Agueci, F., and Vlasenko, L. (2007). Phosphorylation of histone H3 in plants--a dynamic affair. Biochim Biophys Acta 1769, 308–315.

43. Hindriksen, S., Lens, S.M.A., and Hadders, M.A. (2017). The Ins and Outs of Aurora B Inner Centromere Localization. Front Cell Dev Biol 5, 112. 10.3389/fcell.2017.00112.

44. Abad, M.A., Ruppert, J.G., Buzuk, L., Wear, M., Zou, J., Webb, K.M., Kelly, D.A., Voigt, P., Rappsilber, J., Earnshaw, W.C., and Jeyaprakash, A.A. (2019). Borealin-nucleosome interaction secures chromosome association of the chromosomal passenger complex. J Cell Biol 218, 3912–3925. 10.1083/jcb.201905040.

45. Mata-Sucre, Y., Kratka, M., Oliveira, L., Neumann, P., Macas, J., Schubert, V., Huettel, B., Kejnovsky, E., Houben, A., Pedrosa-Harand, A., et al. (2024). Repeat-based holocentromeres of the woodrush Luzula sylvatica reveal insights into the evolutionary transition to holocentricity. Nat Commun 15, 9565. 10.1038/s41467-024-53944-5.

46. Toubiana, W., Dumas, Z., Van, P.T., Parker, D.J., Mérel, V., Schubert, V., Aury, J.-M., Bournonville, L., Cruaud, C., Houben, A., et al. (2025). Functional monocentricity with holocentric characteristics and chromosome-specific centromeres in a stick insect. Science Advances 11, eads6459. 10.1126/sciadv.ads6459.

47. Mirkin, E., and Mirkin, S. (2007). Replication Fork Stalling at Natural Impediments. Microbiology and Molecular Biology Reviews 71, 13–35. 10.1128/mmbr.00030-06.

48. Sproul, J.S., Khost, D.E., Eickbush, D.G., Negm, S., Wei, X., Wong, I., and Larracuente, A.M. (2020). Dynamic Evolution of Euchromatic Satellites on the X Chromosome in Drosophila melanogaster and the simulans Clade. Molecular Biology and Evolution 37, 2241–2256. 10.1093/molbev/msaa078.

49. Kasinathan, S., and Henikoff, S. (2018). Non-B-Form DNA Is Enriched at Centromeres. Molecular biology and evolution 35, 949–962. 10.1093/molbev/msy010.

50. Lermontova, I., Kuhlmann, M., Friedel, S., Rutten, T., Heckmann, S., Sandmann, M., Demidov, D., Schubert, V., and Schubert, I. (2013). Arabidopsis kinetochore null2 is an upstream component for centromeric histone H3 variant cenH3 deposition at centromeres. Plant Cell 25, 3389–3404. 10.1105/tpc.113.114736.

51. Zuo, S., Yadala, R., Yang, F., Talbert, P., Fuchs, J., Schubert, V., Ahmadli, U., Rutten, T., Pecinka, A., Lysak, M.A., and Lermontova, I. (2022). Recurrent Plant-Specific Duplications of KNL2 and Its Conserved Function as a Kinetochore Assembly Factor. Mol Biol Evol 39. 10.1093/molbev/msac123.

52. Hori, T., Shang, W.H., Hara, M., Ariyoshi, M., Arimura, Y., Fujita, R., Kurumizaka, H., and Fukagawa, T. (2017). Association of M18BP1/KNL2 with CENP-A nucleosome is essential for centromere formation in non-mammalian vertebrates. Dev Cell 42, 181–189 e183. 10.1016/j.devcel.2017.06.019.

53. French, B.T., Westhorpe, F.G., Limouse, C., and Straight, A.F. (2017). Xenopus laevis M18BP1 Directly Binds Existing CENP-A Nucleosomes to Promote Centromeric Chromatin Assembly. Dev Cell 42, 190–199 e110. 10.1016/j.devcel.2017.06.021.

54. Maddox, P.S., Hyndman, F., Monen, J., Oegema, K., and Desai, A. (2007). Functional genomics identifies a Myb domain-containing protein family required for assembly of CENP-A chromatin. J Cell Biol 176, 757–763. 10.1083/jcb.200701065.

55. Steiner, F.A., and Henikoff, S. (2014). Holocentromeres are dispersed point centromeres localized at transcription factor hotspots. Elife 3, e02025. 10.7554/eLife.02025.

56. Oliveira, L., Neumann, P., Jang, T.S., Klemme, S., Schubert, V., Koblizkova, A., Houben, A., and Macas, J. (2019). Mitotic spindle attachment to the holocentric chromosomes of Cuscuta europaea does not correlate with the distribution of CENH3 chromatin. Front Plant Sci 10, 1799. 10.3389/fpls.2019.01799.

57. Le Goff, S., Keceli, B.N., Jerabkova, H., Heckmann, S., Rutten, T., Cotterell, S., Schubert, V., Roitinger, E., Mechtler, K., Franklin, F.C.H., et al. (2020). The H3 histone chaperone NASP(SIM3) escorts CenH3 in Arabidopsis. Plant J 101, 71–86. 10.1111/tpj.14518.

58. Takeuchi, H., Nagahara, S., Higashiyama, T., and Berger, F. (2024). The chaperone NASP contributes to de novo deposition of the centromeric histone variant CENH3 in Arabidopsis early embryogenesis. Plant Cell Physiol 65, 1135–1148. 10.1093/pcp/pcae030.

59. Maksimov, V., Nakamura, M., Wildhaber, T., Nanni, P., Ramstrom, M., Bergquist, J., and Hennig, L. (2016). The H3 chaperone function of NASP is conserved in Arabidopsis. Plant J 88, 425–436. 10.1111/tpj.13263.

60. Patthy, L. (2003). Modular assembly of genes and the evolution of new functions. Genetica 118, 217–231.

61. Tromer, E.C., van Hooff, J.J.E., Kops, G., and Snel, B. (2019). Mosaic origin of the eukaryotic kinetochore. Proc Natl Acad Sci U S A 116, 12873–12882. 10.1073/pnas.1821945116.

62. Carmena, M., Wheelock, M., Funabiki, H., and Earnshaw, W.C. (2012). The chromosomal passenger complex (CPC): from easy rider to the godfather of mitosis. Nature Reviews Molecular Cell Biology 13, 789–803. 10.1038/nrm3474.

63. Wang, F., Ulyanova, Natalia P., van der Waal, Maike S., Patnaik, D., Lens, Susanne M.A., and Higgins, Jonathan M.G. (2011). A Positive Feedback Loop Involving Haspin and Aurora B Promotes CPC Accumulation at Centromeres in Mitosis. Current Biology 21, 1061–1069. 10.1016/j.cub.2011.05.016.

64. Venkei, Z., Przewloka, M.R., and Glover, D.M. (2011). Drosophila Mis12 complex acts as a single functional unit essential for anaphase chromosome movement and a robust spindle assembly checkpoint. Genetics 187, 131–140. 10.1534/genetics.110.119628.

65. Ge, X., Ren, J., Gu, K., Gong, W., Shen, K., and Feng, W. (2025). The structure and assembly of the hetero-octameric BLOC-one-related complex. Structure 33, 234–246 e236. 10.1016/j.str.2024.12.001.

66. More, K.J., Kaufman, J.G.G., Dacks, J.B., and Manna, P.T. (2024). Evolutionary origins of the lysosome-related organelle sorting machinery reveal ancient homology in post-endosome trafficking pathways. Proc Natl Acad Sci U S A 121, e2403601121. 10.1073/pnas.2403601121.

67. Nielsen, M., Albrethsen, J., Larsen, F.H., and Skriver, K. (2006). The Arabidopsis ADP-ribosylation factor (ARF) and ARF-like (ARL) system and its regulation by BIG2, a large ARF– GEF. Plant Science 171, 707–717. 10.1016/j.plantsci.2006.07.002.

68. Richardson, D.N., Simmons, M.P., and Reddy, A.S.N. (2006). Comprehensive comparative analysis of kinesins in photosynthetic eukaryotes. Bmc Genomics 7, 18. 10.1186/1471-2164-7-18.

69. Bunning, A.R., and Gupta, M.L., Jr. (2023). The importance of microtubule-dependent tension in accurate chromosome segregation. Front Cell Dev Biol 11, 1096333. 10.3389/fcell.2023.1096333.

70. Dolezel, J., Bartos, J., Voglmayr, H., and Greilhuber, J. (2003). Nuclear DNA content and genome size of trout and human. Cytometry A 51, 127–128; author reply 129. 10.1002/cyto.a.10013.

71. Padmarasu, S., Himmelbach, A., Mascher, M., and Stein, N. (2019). In situ Hi-C for plants: an improved method to detect long-range chromatin interactions. Plant Long Non-Coding RNAs: Methods and Protocols, 441-472.

72. Cheng, H., Concepcion, G.T., Feng, X., Zhang, H., and Li, H. (2021). Haplotype-resolved de novo assembly using phased assembly graphs with hifiasm. Nature methods 18, 170–175. 10.1038/s41592-020-01056-5.

73. Gurevich, A., Saveliev, V., Vyahhi, N., and Tesler, G. (2013). QUAST: quality assessment tool for genome assemblies. Bioinformatics 29, 1072–1075. 10.1093/bioinformatics/btt086.

74. Simão, F.A., Waterhouse, R.M., Ioannidis, P., Kriventseva, E.V., and Zdobnov, E.M. (2015). BUSCO: assessing genome assembly and annotation completeness with single-copy orthologs. Bioinformatics 31, 3210–3212. 10.1093/bioinformatics/btv351.

75. Guan, D., McCarthy, S.A., Wood, J., Howe, K., Wang, Y., and Durbin, R. (2020). Identifying and removing haplotypic duplication in primary genome assemblies. Bioinformatics 36, 2896–2898. 10.1093/bioinformatics/btaa025.

76. Zhou, C., McCarthy, S.A., and Durbin, R. (2023). YaHS: yet another Hi-C scaffolding tool. Bioinformatics 39, btac808.

77. Kim, D., Paggi, J.M., Park, C., Bennett, C., and Salzberg, S.L. (2019). Graph-based genome alignment and genotyping with HISAT2 and HISAT-genotype. Nature biotechnology 37, 907–915.

78. Pertea, M., Pertea, G.M., Antonescu, C.M., Chang, T.-C., Mendell, J.T., and Salzberg, S.L. (2015). StringTie enables improved reconstruction of a transcriptome from RNA-seq reads. Nature biotechnology 33, 290–295.

79. Pertea, G., and Pertea, M. (2020). GFF utilities: GffRead and GffCompare. F1000Research 9.

80. Haas, B.J., Papanicolaou, A., Yassour, M., Grabherr, M., Blood, P.D., Bowden, J., Couger, M.B., Eccles, D., Li, B., and Lieber, M. (2013). De novo transcript sequence reconstruction from RNA-seq using the Trinity platform for reference generation and analysis. Nature protocols 8, 1494–1512.

81. Gabriel, L., Bruna, T., Hoff, K.J., Ebel, M., Lomsadze, A., Borodovsky, M., and Stanke, M. (2024). BRAKER3: Fully automated genome annotation using RNA-seq and protein evidence with GeneMark-ETP, AUGUSTUS, and TSEBRA. Genome Res 34, 769–777. 10.1101/gr.278090.123.

82. He, W., Yang, J., Jing, Y., Xu, L., Yu, K., and Fang, X. (2023). NGenomeSyn: an easy-to-use and flexible tool for publication-ready visualization of syntenic relationships across multiple genomes. Bioinformatics 39, btad121.

83. Grabherr, M.G., Haas, B.J., Yassour, M., Levin, J.Z., Thompson, D.A., Amit, I., Adiconis, X., Fan, L., Raychowdhury, R., and Zeng, Q. (2011). Full-length transcriptome assembly from RNA-Seq data without a reference genome. Nature biotechnology 29, 644–652.

84. Komaki, S., and Schnittger, A. (2017). The Spindle Assembly Checkpoint in Arabidopsis Is Rapidly Shut Off during Severe Stress. Dev Cell 43, 172–185 e175. 10.1016/j.devcel.2017.09.017.

85. Su, H., Liu, Y., Wang, C., Liu, Y., Feng, C., Sun, Y., Yuan, J., Birchler, J.A., and Han, F. (2021). Knl1 participates in spindle assembly checkpoint signaling in maize. Proc Natl Acad Sci U S A 118. 10.1073/pnas.2022357118.

86. Rice, P., Longden, I., and Bleasby, A. (2000). EMBOSS: the European Molecular Biology Open Software Suite. Trends Genet 16, 276–277. 10.1016/s0168-9525(00)02024-2.

87. Birney, E., Clamp, M., and Durbin, R. (2004). GeneWise and Genomewise. Genome Res 14, 988–995. 10.1101/gr.1865504.

88. Thompson, J.D., Higgins, D.G., and Gibson, T.J. (1994). Clustal-W - Improving the Sensitivity of Progressive Multiple Sequence Alignment through Sequence Weighting, Position-Specific Gap Penalties and Weight Matrix Choice. Nucleic Acids Research 22, 4673–4680. DOI 10.1093/nar/22.22.4673.

89. Kumar, S., Stecher, G., Li, M., Knyaz, C., and Tamura, K. (2018). MEGA X: Molecular Evolutionary Genetics Analysis across Computing Platforms. Molecular Biology and Evolution 35, 1547–1549. 10.1093/molbev/msy096.

90. Trifinopoulos, J., Nguyen, L.T., von Haeseler, A., and Minh, B.Q. (2016). W-IQ-TREE: a fast online phylogenetic tool for maximum likelihood analysis. Nucleic Acids Research 44, W232–W235. 10.1093/nar/gkw256.

91. Letunic, I., and Bork, P. (2007). Interactive Tree Of Life (iTOL): an online tool for phylogenetic tree display and annotation. Bioinformatics 23, 127–128. 10.1093/bioinformatics/btl529.

92. Letunic, I., and Bork, P. (2019). Interactive Tree Of Life (iTOL) v4: recent updates and new developments. Nucleic Acids Research 47, W256–W259. 10.1093/nar/gkz239.

93. Dolezel, J., Binarova, P., and Lucretti, S. (1989). Analysis of nuclear DNA content in plant cells by flow cytometry. Biologia Plantarum 31, 113–120.

94. Pellicer, J., Fernandez, P., Fay, M.F., Michalkova, E., and Leitch, I.J. (2021). Genome Size Doubling Arises From the Differential Repetitive DNA Dynamics in the Genus Heloniopsis (Melanthiaceae). Front Genet 12, 726211. 10.3389/fgene.2021.726211.

95. Andrews, S. (2010). FastQC: a quality control tool for high throughput sequence data.

96. Novak, P., Neumann, P., and Macas, J. (2010). Graph-based clustering and characterization of repetitive sequences in next-generation sequencing data. BMC Bioinformatics 11, 378. 10.1186/1471-2105-11-378.

97. Novak, P., Neumann, P., Pech, J., Steinhaisl, J., and Macas, J. (2013). RepeatExplorer: a Galaxy-based web server for genome-wide characterization of eukaryotic repetitive elements from next-generation sequence reads. Bioinformatics 29, 792–793. 10.1093/bioinformatics/btt054.

98. Novák, P., Neumann, P., and Macas, J. (2020). Global analysis of repetitive DNA from unassembled sequence reads using RepeatExplorer2. Nature Protocols 15, 3745–3776. 10.1038/s41596-020-0400-y.

99. Bolger, A.M., Lohse, M., and Usadel, B. (2014). Trimmomatic: a flexible trimmer for Illumina sequence data. Bioinformatics 30, 2114–2120. 10.1093/bioinformatics/btu170.

100. Langmead, B., and Salzberg, S.L. (2012). Fast gapped-read alignment with Bowtie 2. Nature methods 9, 357–359.

101. Ramirez, F., Ryan, D.P., Gruning, B., Bhardwaj, V., Kilpert, F., Richter, A.S., Heyne, S., Dundar, F., and Manke, T. (2016). deepTools2: a next generation web server for deep-sequencing data analysis. Nucleic Acids Res 44, W160–165. 10.1093/nar/gkw257.

102. Lopez-Delisle, L., Rabbani, L., Wolff, J., Bhardwaj, V., Backofen, R., Grüning, B., Ramírez, F., and Manke, T. (2021). pyGenomeTracks: reproducible plots for multivariate genomic data sets. Bioinformatics.

103. Vollger, M.R., Kerpedjiev, P., Phillippy, A.M., and Eichler, E.E. (2022). StainedGlass: interactive visualization of massive tandem repeat structures with identity heatmaps. Bioinformatics 38, 2049–2051. 10.1093/bioinformatics/btac018.

104. Chen, S., Zhou, Y., Chen, Y., and Gu, J. (2018). fastp: an ultra-fast all-in-one FASTQ preprocessor. Bioinformatics 34, i884–i890. 10.1093/bioinformatics/bty560.

105. Vasimuddin, M., Misra, S., Li, H., and Aluru, S. (2019). Efficient architecture-aware acceleration of BWA-MEM for multicore systems. (IEEE), pp. 314–324.

106. Li, H., Handsaker, B., Wysoker, A., Fennell, T., Ruan, J., Homer, N., Marth, G., Abecasis, G., Durbin, R., and Genome Project Data Processing, S. (2009). The Sequence Alignment/Map format and SAMtools. Bioinformatics 25, 2078–2079. 10.1093/bioinformatics/btp352.

107. Zhang, Y., Liu, T., Meyer, C.A., Eeckhoute, J., Johnson, D.S., Bernstein, B.E., Nusbaum, C., Myers, R.M., Brown, M., Li, W., and Liu, X.S. (2008). Model-based Analysis of ChIP-Seq (MACS). Genome Biology 9, R137. 10.1186/gb-2008-9-9-r137.

108. Novak, P., Avila Robledillo, L., Koblizkova, A., Vrbova, I., Neumann, P., and Macas, J. (2017). TAREAN: a computational tool for identification and characterization of satellite DNA from unassembled short reads. Nucleic Acids Res 45, e111. 10.1093/nar/gkx257.

109. Richards, E.J., and Ausubel, F.M. (1988). Isolation of a higher eukaryotic telomere from Arabidopsis thaliana. Cell 53, 127–136. 10.1016/0092-8674(88)90494-1.

110. Zhang, T., Liu, G., Zhao, H., Braz, G.T., and Jiang, J. (2021). Chorus2: design of genome-scale oligonucleotide-based probes for fluorescence in situ hybridization. Plant Biotechnol J 19, 1967–1978. 10.1111/pbi.13610.

111. Baez, M., Kuo, Y.T., Dias, Y., Souza, T., Boudichevskaia, A., Fuchs, J., Schubert, V., Vanzela, A.L.L., Pedrosa-Harand, A., and Houben, A. (2020). Analysis of the small chromosomal Prionium serratum (Cyperid) demonstrates the importance of reliable methods to differentiate between mono- and holocentricity. Chromosoma 129, 285–297. 10.1007/s00412-020-00745-6.

112. Weisshart, K., Fuchs, J., and Schubert, V. (2016). Structured illumination microscopy (SIM) and photoactivated localization microscopy (PALM) to analyze the abundance and distribution of RNA polymerase II molecules on flow-sorted Arabidopsis nuclei. Bio-protocol 6, e1725–e1725.

113. Daghma, D.S., Kumlehn, J., and Melzer, M. (2011). The use of cyanobacteria as filler in nitrocellulose capillaries improves ultrastructural preservation of immature barley pollen upon high pressure freezing. J Microsc-Oxford 244, 79–84. 10.1111/j.1365-2818.2011.03509.x.

## Supplemental references

1. Pellicer, J., Fernandez, P., Fay, M.F., Michalkova, E., and Leitch, I.J. (2021). Genome Size Doubling Arises From the Differential Repetitive DNA Dynamics in the Genus Heloniopsis (Melanthiaceae). Front Genet 12, 726211. 10.3389/fgene.2021.726211.

2. Kuo, Y.-T., Câmara, A.S., Schubert, V., Neumann, P., Macas, J., Melzer, M., Chen, J., Fuchs, J., Abel, S., Klocke, E., et al. (2023). Holocentromeres can consist of merely a few megabase-sized satellite arrays. Nat Commun 14, 3502. 10.1038/s41467-023-38922-7.

## Supplemental references

1. Naish, M., Alonge, M., Wlodzimierz, P., Tock, A.J., Abramson, B.W., Schmücker, A., Mandáková, T., Jamge, B., Lambing, C., Kuo, P., et al. (2021). The genetic and epigenetic landscape of the Arabidopsis centromeres. Science 374. 10.1126/science.abi7489.

2. Chen, C., Wu, S., Sun, Y., Zhou, J., Chen, Y., Zhang, J., Birchler, J.A., Han, F., Yang, N., and Su, H. (2024). Three near-complete genome assemblies reveal substantial centromere dynamics from diploid to tetraploid in Brachypodium genus. Genome Biol 25, 63. 10.1186/s13059-024-03206-w.

3. Zhang, L., Liang, J., Chen, H., Zhang, Z., Wu, J., and Wang, X. (2023). A near-complete genome assembly of Brassica rapa provides new insights into the evolution of centromeres. Plant Biotechnol J 21, 1022–1032. 10.1111/pbi.14015.

4. Chen, W., Wang, X., Sun, J., Wang, X., Zhu, Z., Ayhan, D.H., Yi, S., Yan, M., Zhang, L., Meng, T., et al. (2024). Two telomere-to-telomere gapless genomes reveal insights into Capsicum evolution and capsaicinoid biosynthesis. Nat Commun 15, 4295. 10.1038/s41467-024-48643-0.

5. Wang, B., Jia, Y., Dang, N., Yu, J., Bush, S.J., Gao, S., He, W., Wang, S., Guo, H., Yang, X., et al. (2024). Near telomere-to-telomere genome assemblies of two Chlorella species unveil the composition and evolution of centromeres in green algae. Bmc Genomics 25, 356. 10.1186/s12864-024-10280-8.

6. Xia, Q.M., Miao, L.K., Xie, K.D., Yin, Z.P., Wu, X.M., Chen, C.L., Grosser, J.W., and Guo, W.W. (2020). Localization and characterization of Citrus centromeres by combining half-tetrad analysis and CenH3-associated sequence profiling. Plant Cell Rep 39, 1609–1622. 10.1007/s00299-020-02587-z.

7. Chang, C.H., Chavan, A., Palladino, J., Wei, X., Martins, N.M.C., Santinello, B., Chen, C.C., Erceg, J., Beliveau, B.J., Wu, C.T., et al. (2019). Islands of retroelements are major components of Drosophila centromeres. PLoS Biol 17, e3000241. 10.1371/journal.pbio.3000241.

8. Cui, J., Zhu, C., Shen, L., Yi, C., Wu, R., Sun, X., Han, F., Li, Y., and Liu, Y. (2024). The gap-free genome of Forsythia suspensa illuminates the intricate landscape of centromeres. Hortic Res 11, uhae185. 10.1093/hr/uhae185.

9. Liu, Y., Yi, C., Fan, C., Liu, Q., Liu, S., Shen, L., Zhang, K., Huang, Y., Liu, C., Wang, Y., et al. (2023). Pan-centromere reveals widespread centromere repositioning of soybean genomes. Proc Natl Acad Sci U S A 120, e2310177120. 10.1073/pnas.2310177120.

10. Huang, G., Bao, Z., Feng, L., Zhai, J., Wendel, J.F., Cao, X., and Zhu, Y. (2024). A telomere-to-telomere cotton genome assembly reveals centromere evolution and a Mutator transposon-linked module regulating embryo development. Nat Genet 56, 1953–1963. 10.1038/s41588-024-01877-6.

11. Ding, W., Zhu, Y., Han, J., Zhang, H., Xu, Z., Khurshid, H., Liu, F., Hasterok, R., Shen, X., and Wang, K. (2023). Characterization of centromeric DNA of Gossypium anomalum reveals sequence-independent enrichment dynamics of centromeric repeats. Chromosome Res 31, 12. 10.1007/s10577-023-09721-z.

12. Logsdon, G.A., Rozanski, A.N., Ryabov, F., Potapova, T., Shepelev, V.A., Catacchio, C.R., Porubsky, D., Mao, Y., Yoo, D., Rautiainen, M., et al. (2024). The variation and evolution of complete human centromeres. Nature 629, 136–145. 10.1038/s41586-024-07278-3.

13. Dias, Y., Mata-Sucre, Y., Thangavel, G., Costa, L., Báez, M., Houben, A., Marques, A., and Pedrosa-Harand, A. (2024). How diverse a monocentric chromosome can be? Repeatome and centromeric organization of Juncus effusus (Juncaceae). The Plant Journal n/a. 10.1111/tpj.16712.

14. Wang, K., Jin, J., Wang, J., Wang, X., Sun, J., Meng, D., Wang, X., Wang, Y., and Guo, L. (2024). The complete telomere-to-telomere genome assembly of lettuce. Plant Commun 5, 101011. 10.1016/j.xplc.2024.101011.

15. Chen, W., Yan, M., Chen, S., Sun, J., Wang, J., Meng, D., Li, J., Zhang, L., and Guo, L. (2024). The complete genome assembly of Nicotiana benthamiana reveals the genetic and epigenetic landscape of centromeres. Nat Plants 10, 1928–1943. 10.1038/s41477-024-01849-y.

16. Song, J.M., Xie, W.Z., Wang, S., Guo, Y.X., Koo, D.H., Kudrna, D., Gong, C., Huang, Y., Feng, J.W., Zhang, W., et al. (2021). Two gap-free reference genomes and a global view of the centromere architecture in rice. Mol Plant 14, 1757–1767. 10.1016/j.molp.2021.06.018.

17. Liu, C., Fu, S., Yi, C., Liu, Y., Huang, Y., Guo, X., Zhang, K., Liu, Q., Birchler, J.A., and Han, F. (2024). Unveiling the distinctive traits of functional rye centromeres: minisatellites, retrotransposons, and R-loop formation. Sci China Life Sci 67, 1989–2002. 10.1007/s11427-023-2524-0.

18. Pham, G.M., Hamilton, J.P., Wood, J.C., Burke, J.T., Zhao, H., Vaillancourt, B., Ou, S., Jiang, J., and Buell, C.R. (2020). Construction of a chromosome-scale long-read reference genome assembly for potato. Gigascience 9. 10.1093/gigascience/giaa100.

19. Ahmed, H.I., Heuberger, M., Schoen, A., Koo, D.H., Quiroz-Chavez, J., Adhikari, L., Raupp, J., Cauet, S., Rodde, N., Cravero, C., et al. (2023). Einkorn genomics sheds light on history of the oldest domesticated wheat. Nature 620, 830–838. 10.1038/s41586-023-06389-7.

20. Su, H., Liu, Y., Liu, C., Shi, Q., Huang, Y., and Han, F. (2019). Centromere Satellite Repeats Have Undergone Rapid Changes in Polyploid Wheat Subgenomes. Plant Cell 31, 2035–2051. 10.1105/tpc.19.00133.

21. Shi, X., Cao, S., Wang, X., Huang, S., Wang, Y., Liu, Z., Liu, W., Leng, X., Peng, Y., Wang, N., et al. (2023). The complete reference genome for grapevine (Vitis vinifera L.) genetics and breeding. Hortic Res 10, uhad061. 10.1093/hr/uhad061.

22. Chen, J., Wang, Z., Tan, K., Huang, W., Shi, J., Li, T., Hu, J., Wang, K., Wang, C., Xin, B., et al. (2023). A complete telomere-to-telomere assembly of the maize genome. Nat Genet 55, 1221–1231. 10.1038/s41588-023-01419-6.

23. Kuo, Y.-T., Câmara, A.S., Schubert, V., Neumann, P., Macas, J., Melzer, M., Chen, J., Fuchs, J., Abel, S., Klocke, E., et al. (2023). Holocentromeres can consist of merely a few megabase-sized satellite arrays. Nat Commun 14, 3502. 10.1038/s41467-023-38922-7.

